# Converting *Escherichia coli* MG1655 into a chemical overproducer through inactivating defense system against exogenous DNA

**DOI:** 10.1101/2020.08.15.251348

**Authors:** Jingge Wang, Chaoyong Huang, Kai Guo, Lianjie Ma, Xiangyu Meng, Ning Wang, Yi-Xin Huo

**Affiliations:** Key Laboratory of Molecular Medicine and Biotherapy, School of Life Sciences, Beijing Institute of Technology, No. 5 South Zhongguancun Street, Beijing, China, 100081; SIP-UCLA Institute for Technology Advancement, 10 Yueliangwan Road, Suzhou Industrial Park, Suzhou, China, 215123; Biology Institute, Shandong Province Key Laboratory for Biosensors, Qilu University of Technology (Shandong Academy of Sciences), Jinan, 250103 China

**Keywords:** *Escherichia coli*, MG1655, CRISPR‐Cas9, defense system, metabolic engineering

## Abstract

*Escherichia coli* strain K-12 MG1655 has been proposed as an appropriate host strain for industrial production. However, the direct application of this strain suffers from the transformation inefficiency and plasmid instability. Herein, we conducted genetic modifications at a serial of loci of MG1655 genome, generating a robust and universal host strain JW128 with higher transformation efficiency and plasmid stability that can be used to efficiently produce desired chemicals after introducing the corresponding synthetic pathways. Using JW128 as the host, the titer of isobutanol reached 5.76 g/L in shake-flask fermentation, and the titer of lycopene reached 1.91 g/L in test-tube fermentation, 40-fold and 5-fold higher than that of original MG1655, respectively. These results demonstrated JW128 is a promising chassis for high-level production of value-added chemicals.

## 1. Introduction

*Escherichia coli*, as one of the most important microorganism hosts, has been widely used for producing value-added chemicals [1–3]. Up to now, many *E. coli* strains have served as the industrial hosts, including BL21, C, Crooks, DH5α, K-12 MG1655, K-12 W3110, and W [4–10]. By using genomics, phenomics, transcriptomics, and genome-scale modeling, *Monk et al*. computationally predicted that K-12 MG1655 has the greatest producing potential among the seven *E. coli* strains for two types of compounds under aerobic and anaerobic conditions: (1) all 20 amino acids using native pathways and (2) non-native compounds using 245 heterologous pathways [4]. Combining predicted fluxes with gene expression values would generate a relative production potential score (‘‘R score’’) that gauged a strain’s suitability for producing a given compound. Under both aerobic and anaerobic conditions, MG1655 frequently had an R score >1 for amino acid production (12/20 and 7/20, respectively), whereas K-12 MG1655 had the highest number of R scores >1 for 94 aerobic pathways during the production potential examination across 245 heterologous pathways (corresponding to one of 20 diverse industrial compounds, originated from cell native precursors) [4]. Besides an optimal metabolic network, high growth rate and strong robustness endow MG1655 a great potential to be applied in industrial production. Although MG1655 has many advantages as a metabolic engineering host, its large-scale application is limited by some intrinsic properties.

MG1655 was chosen by the Blattner group for the first published sequence of *E. coli*, which is a wild-type laboratory strain that has few genetic manipulations from the archetypal *E. coli* K-12 strain [11]. Like other wild-type *E. coli* strains, MG1655 has poorer competency and plasmid stability compared with commercial *E. coli* strains, which influence the ease and labor of genomic editing for constructing fermentation strains. For example, DH10B is one of the commercial strains that is frequently applied as a host strain for molecular cloning due to its incomplete defense system for exogenous DNA [12]. The MG1655 defense system mainly consists of DNA-specific endonuclease I EndA and the restriction modification system (R-M system). EndA is a periplasmic enzyme that cleaves double-strand DNA (dsDNA), thereby, affecting the stability of the plasmid DNA [13]. A classic R-M system includes an endonuclease that cleaves a specific DNA sequence and a DNA methyltransferase that methylates either adenosyl or cytosyl residues within the same DNA sequence [14–16]. The R-M system consists of seven genes including *hsdRMS* [16], *mcrA* [17, 18], *mcrBC* [19, 20], and *mrr* [21]. Apart from these, *recA* gene also contribute to the instability of plasmid DNA. Inactivation of genes of the defense system will greatly increase the plasmid DNA stability in MG1655, resulting in the increase of transformation efficiency and biomolecule productivity. Hence, an appropriate genetic manipulation tool is required to inactivate genes precisely and efficiently.

During the last decades, a variety of genome editing methods have been developed, and the most popular ones are based on homologous recombination. A classic and commonly used method is recombineering that utilizes bacteriophage-derived λ-Red system to enhance the recombination activities of *E. coli* [22–24]. Though recombineering can handle simple DNA manipulations [25–27], its editing efficiency depends on the sequence length [22]. Moreover, eliminating selectable markers and plasmids is complicated and time-consuming, and the residual *FRT* or *loxP* site may influence the next round of genome editing [28]. Prior to homologous recombination, generating a double-strand break (DSB) in the target DNA is an effective strategy for improving the editing efficiency of complex genetic manipulations and avoiding the introduction of selectable markers and specific recombination sites. Compared with methods that utilize homing endonuclease *I-SceI* [29–31] and engineered endonucleases, such as zinc-finger nucleases (ZFNs) and transcription activator-like effector nucleases (TALENs) [32–35], to cleave dsDNA, clustered regularly interspaced short palindromic repeats (CRISPR)/CRISPR-associated protein 9 (Cas9) system is a more efficient and convenient route to generate DSB [36–38]. Taken together, the combination of recombineering and CRISPR/Cas9 technologies is a better strategy for gene knockouts.

In this study, we demonstrated advantages of MG1655 in growth rate and robustness by comparing it with several commonly used commercial *E. coli* strains. Genes involved in the defense system against foreign DNA were inactivated, generating a robust and universal chassis strain for metabolic engineering. The engineered strain not only reserved all merits of MG1655, but also acquired new merits including superior competency and plasmid stability. We also demonstrated the potential of the strain in metabolic engineering applications by using it as a host to produce isobutanol and lycopene.

## 2. Materials and methods

### 2.1. Strains, plasmids, reagents and growth conditions

The plasmids and strains used in this study, as well as their relevant characteristics, were shown in Supplementary Table 3 and Table 4. *E. coli* DH5α was used as a plasmid cloning host strain and MG1655 was used as the chassis strain. Luria-Bertani (LB) medium was used for cell growth in all cases unless otherwise noted. Agar was added at 20 g/L for LB solid medium. The M9Y medium was used for the production of isobutanol. SOC medium was used for cell recovery in the transformation process and genome editing procedure. LB, SOB, and SOC mediums were also used in the process of the optimization of making competent cells. Where appropriate, ampicillin (Amp), kanamycin (Kan), IPTG, _L_-arabinose, glucose was added. The reagents and mediums used in this study were shown in supplementary Table 1.

### 2.2. Determination of growth curves and robustness of *E. coli* strains via OD measurement

Eight strains MG1655, DH5α, DH10B, TOP10, BL21(DE3), XL1-blue, XL10-gold, and Mach1-T1 were tested in this study. For the preculture, a colony was picked and cultured overnight in 5 mL LB at 37 °C, and 1 mL overnight culture was added to 100 mL LB medium in 250 mL flask. The initial OD_600_, of each strain in the main culture should be consistent.

For the measurement of growth curves, these strains grew at 37 °C with shaking for 12 hours. During the 12 hours, samples were taken every hour and stored at 4 °C. The optical density of the samples was measured by using a spectrophotometer (METASH, V-5100) at 600 nm. For the robustness measurement, these strains were cultured under three types of stressed conditions for 12 hours and the OD_600_ of eight strains were measured. The three types of stressed factors are temperature, acid, and salinity.

### 2.3. Preparation of chemical and electroporation competent cells

Competent cells were prepared via two methods: the classic CaCl_2_ and electroporation methods. The electrocompetent cells were used for genome editing. The chemical competent cells were used for plasmid construction and transformation efficiency measurement.

The plasmid p-P_BAD_-*cas9*/P_T5_-Redγβα was transformed into *E. coli* MG1655 and the transformant was made into electrocompetent cells for genome editing. A colony was cultured overnight in 5 mL LB at 37 °C, then 0.5 mL overnight medium and 1 mL glucose were transferred into 50 mL LB medium. The medium was cultured at 37 °C, 220 rpm until the OD_600_ reached 0.4-0.6. After 30 minutes in the ice bath, the bacterial solution was pelleted by centrifugation at 1520 × g for 10 min at 4 °C. The cells were washed twice with 10 mL ice-cold 10% glycerol and resuspended with 1.5 mL ice-cold 10% glycerol. Then the cells were divided into 100-μl aliquots and stored at −80 °C. For making chemical competent cells, glucose was needless and the operations before wash the cells were the same as described above. The chemical competent cells were washed twice with ice-cold CaCl_2_ (0.1 M) and resuspended in the mixture of 0.9 mL CaCl_2_ and 0.6 mL 50% glycerol. The competent cells were also divided into 100-μl aliquots and stored at −80 °C.

### 2.4. Iterative genome editing procedure

The genome editing process mainly included three steps. Firstly, the Kan‐resistant plasmid p-P_BAD_-*cas9*/P_T5_-Redγβα was transferred into the MG1655 to obtain the transformant MG1655/p-P_BAD_-*cas9*/P_T5_-Redγβα. Secondly, the temperature-sensitive Amp‐resistant plasmid p-PBAD-sgRNA-X was transformed into the strain MG1655/p-P_BAD_-*cas9*/P_T5_-Redγβα, the recombinant strain MG1655/p-P_BAD_-*cas9*/P_T5_-Redγβα/p-P_BAD_-sgRNA-X was cultured on LB plate carrying Amp, Kan, and glucose at 30 °C. A single colony was picked and cultured in 0.5 mL SOC at 30 °C. After 2 hours of cultivation, 4.5 mL LB, 5 μl Amp, 5 μl Kan, and 50 μl IPTG were added to the cultures. After 1 h, 100 μl _L-_arabinose was added to the cultures and incubated for another 3 h. A 1 μl aliquot of the cultures was added to the 100 μl sterile water and cultured on LB plate containing Amp, Kan, and _L_-arabinose overnight at 30 °C. The colonies were analyzed by the colony polymerase chain reaction (PCR). The positive PCR products were subject to verification by DNA sequencing for further confirmation. After one round of editing, the strain was cultured overnight in LB supplemented with Kan at 37 °C to cure the temperature-sensitive plasmid p-P_BAD_-sgRNA-X. The strain harboring plasmid p-P_BAD_-*cas9*/P_T5_-Redγβα was made into competent cells and another editing plasmid p-PBAD-sgRNA-X was transformed into the competent cells for the next round of editing. The primers used in this study were shown in supplementary Table 5.

### 2.5. Plasmid curing

After five rounds of genome editing, in order to cure the plasmid, the strain harboring Kan^R^ plasmid p-P_BAD_-*cas9*/P_T5_-Redγβα was cultured in LB at 37 °C for 16 hours with no antibiotics. The culture was diluted and plated onto an LB plate with no antibiotics and then incubated overnight at 37 °C. The colonies were randomly picked and transferred on LB plates containing Kan and no antibiotics, respectively, and cultured at 37 °C. The colonies lacking plasmid cannot survive in the culture with Kan. The plasmid-free strains were used in other experiments in this study, such as transformation efficiency, growth curve measurement, and chemical production.

### 2.6. Measurement of the transformation efficiency

Plasmid pUC19 was used to measure the transformation efficiency of chemical competent cells. One tube of a competent cell (100 μl) from −80 °C was thawed on ice. A total of 10 ng pure pUC19 plasmid DNA was added to cells. The tube was stored on ice for 30 min, heated at 42 °C for 1 min, and then immediately transferred on ice for 2 min. 1mL SOC medium was added into the tube and incubated with gentle shaking (220 rpm) at 37 °C for 40 min. 100 μl of the cell suspension cultures were plated on LB plate with Amp and cultured overnight at 37 °C. The transformation efficiency was N × 10^3^ cfu/μg DNA. (“N” refers to the total colony forming units).

### 2.7. The fermentation of higher alcohol

Two pairs of combinational plasmids were used to produce isobutanol, one was pSA65 and pSA69, the other one was pSA69 and pYX97. The plasmids pSA69 and pSA65 were transferred into the experimental strains MG1655, JW128, and JCL16. To prepare the seed culture, one colony containing two plasmids was cultured in 5 mL LB with 5 μl Amp and 5 μl Kan at 37 °C and 220 rpm overnight. The seed culture was inoculated into a 250 mL shake flask with 20 mL M9Y medium, 0.1 mM IPTG, 0.1 g/L Amp, 0.05 g/L Kan, and incubated at 30 °C and 250 rpm. The inoculation ratio of seed culture was 1/100, the other strains were adjusted according to the OD_600_ value of seed culture to make the initial OD_600_ consistent. The plasmids pSA69 and pYX97 were transferred into the strains JW128, and JCL260, the other operations were the same as mentioned above. The samples were taken every 12 h and the biomass concentration was evaluated by using a spectrometer (METASH, V-5100) at 600 nm. The remaining samples were centrifuged at 16,060 ×g for 2 min, and the supernatants were stored at-20 °C for the determination of isobutanol content.

### 2.8. Gas chromatography (GC) detection of higher alcohol

Production of isobutanol was quantified by Agilent 6890 GC chromatograph equipped with a flame ionization detector (Agilent Technologies, CA, USA). Nitrogen was used as a carrier gas. The GC oven temperature was initially at 80 °C for 3 min, and increased to 230 °C at a rate of 115 °C/min, and held for 1 min. The injector and detector were maintained at 250 °C and 280 °C, respectively. The supernatant (1 μl) was sampled and injected at a split ratio of 1:30 and pentanol was used as an internal standard [39]. The internal standard method was used to measure the production of biofuels. All experiments were done in a triple, and data were expressed as the mean ± standard error by using GraphPad prism.

### 2.9. The production of lycopene

The plasmid p‐P_fumAp_‐*crtEBI* was transformed into four strains MG1655, JW128, DH10B, and DH5α to produce lycopene. To make the seed culture, a colony was picked and cultured overnight in 5 mL LB containing 5 μl Amp at 37 °C and 220 rpm. Then 50 μl seed culture was inoculated into 5 mL LB with 5 μl Amp and cultured at 37 °C, 220 rpm for 22 hours. The volume of the seed culture of other strains was adjusted according to the OD_600_ value to make the initial bacterial amount as consistent as possible. After 22 hours, the OD_600_ of four strains were measured and 4 mL bacterial cultures were pelleted at 16,060×g for 3 min. Cell pellets were resuspended in 1 mL ultrapure water and centrifuged again to remove the supernatant. 1 mL acetone was added to the cell and the mixture of cell and acetone was placed at 56 °C for 20 min. Then the cell mixture was centrifuged at 16,060 ×g for 3 min. 500 μl of supernatant was removed for analysis by using a spectrometer (METASH, V-5100) at 474 nm [40], 3 mL ultrapure water was used as an internal standard. The standard lycopene (CAS:502-65-8) was used to measure the standard curve. All experiments were done in a triple, and data were expressed as the mean ± standard error by using GraphPad prism.

## 3. Results and discussion

### 3.1. Growth advantage of MG1655 under unstressed and stressed conditions

To investigate the comparative growth of MG1655 with its sibling *E. coli* strains, seven common commercial *E. coli* strains were selected as the control group. The OD_600_ values of eight strains were measured under all conditions and the growth curves were plotted by the Growth fitting. The unstressed growth condition was set as the 37°C in LB culture, while the stressed condition groups included heat-, acid-, and osmotic shocks which frequently occur during the routine fermentations.

At 12 h, the OD_600_ of MG1655 under unstressed growth condition reached 5.49 while the one of DH5α, DH10B, TOP10, BL21(DE3), XL1-blue, XL10-gold, and Mach1‐T1 was 3.64, 2.99, 3.20, 5.01, 4.64, 3.78, and 3.85, respectively. Based on the growth curve (Fig. 1a and Fig. S1), the specific growth rate (h^−1^) of each strain was calculated (Fig. 1a). Under unstressed growth condition, the specific growth rate of MG1655 was 1.40 h^−1^ while the one of other strains varied from 1.02 h^−1^ to 1.38 h^−1^. By comparison of the values of OD_600_ and the specific growth rate, MG1655 showed the biomass and growth rate advantages under unstressed condition. To represent the robustness, the rangeability of OD_600_ of eight strains were calculated when the strains were cultured under the three stressed conditions (Fig. 1b). Among the three stress factors, osmotic stress had the strongest impact for all strains, while the acid stress had the weakest impact for all strains. Under osmotic-shock condition, the relative changes of OD_600_ of MG1655 was 15.8% while the one of DH5α, DH10B, TOP10, BL21(DE3), XL1-blue, XL10-gold, and Mach1-T1 was 39.3%, 92.4%, 26.9%, 39.9%, 44.6%, 66.8%, and 42.1%, respectively. Under acid-shock condition, the OD_600_ relative changes of MG1655 was 0.5%, while the one of other strains ranged from 2.5% to 21.4%. During the heat-shock condition, the OD_600_ relative changes of MG1655 were 2.6%, while the one of other strains were between 5.2% and 27%. The maximum tolerability of each strain under osmotic-, acid- and heat-stressed conditions were also measured (Fig. S2). The growth of eight strains weakened with the increase of environmental stresses. Under osmotic-stressed conditions with 60 g/L NaCl, MG1655 was least affected by the osmotic stress and the OD_600_ reached 1.42, while the one of other strains ranged from 0.1 to 0.46 (Fig. S2a). Under acid-stressed conditions, the maximum pH tolerability of XL10-gold was 4.5, the other seven strains could survive in the medium of pH 4. Moreover, the OD_600_ value of MG1655 was highest, it reached 1.77, while that of other six strains ranged from 1.04 to 1.33 (Fig. S2b). Under heat-stressed conditions, the maximum tolerability of eight strains were 45 °C. However, the OD_600_ value of MG1655 was highest and it reached 3.92, while the one of other strains were between 2.00 and 3.53 (Fig. S2c).

**Fig. 1.**
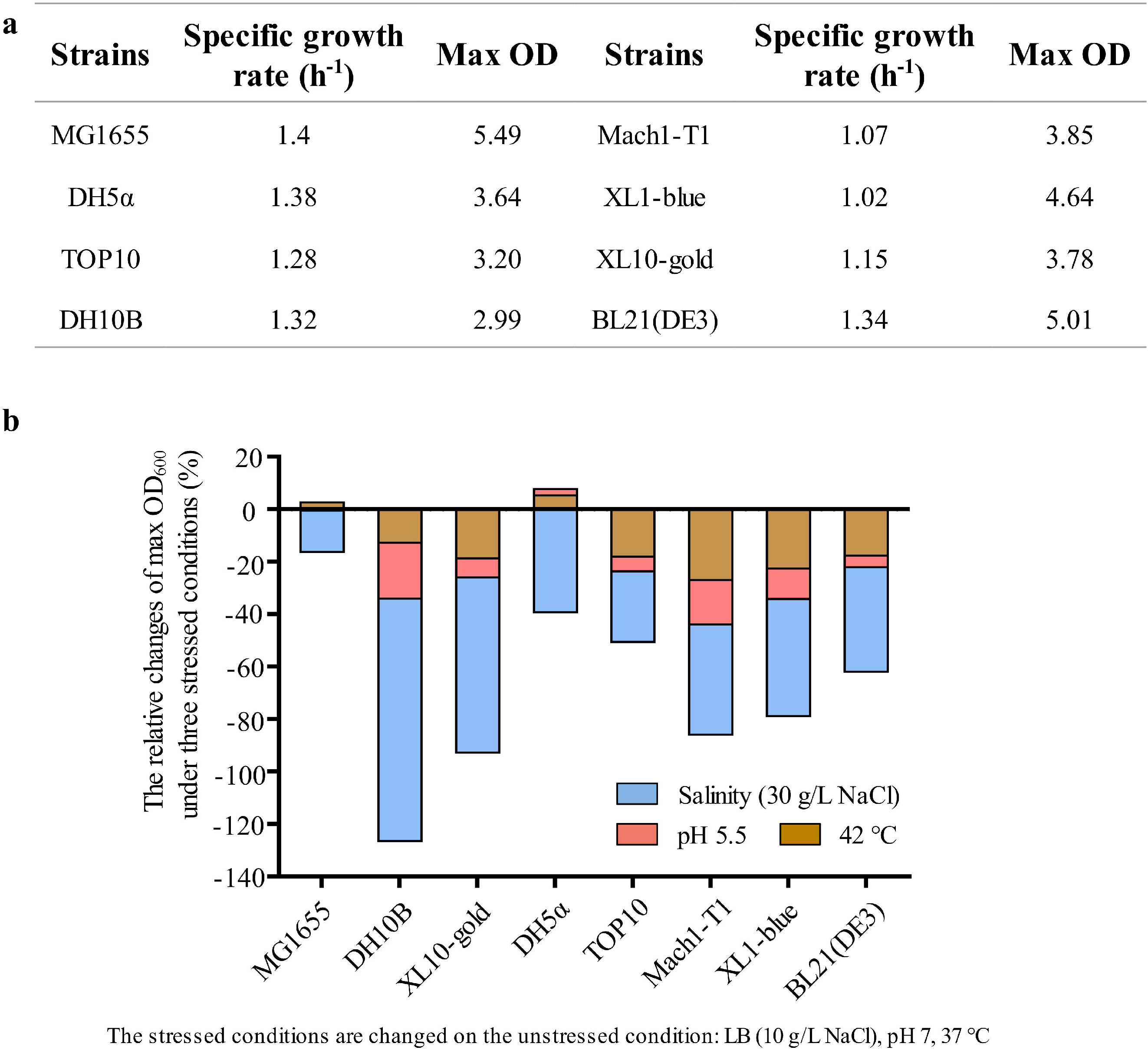
The comparison of MG1655 and seven *E. coli* strains in growth rate and robustness. (a) The detailed number of specific growth rate (h^−1^) and max OD of the eight strains cultured in the unstressed condition (LB, pH 7, 37 °C). (b) The fluctuation of max OD of eight strains under three stressed conditions compared to the unstressed condition. Three stressed conditions changed the culture temperature, pH, and the salinity of LB medium, respectively. The relative changes = (max OD _unstressed_ – max OD _stressed_) / max OD _unstressed_ * 100%.

Taken together, MG1655 presented the highest growth rate comparing with its seven sibling *E. coli* strains. Generally, a higher growth rate will endow a stronger biomass building capacity of the strain, therefore efficiently promoting the strain to convert available carbon source into key metabolic intermediates. Since the key metabolic intermediates could be hijacked into bulk chemicals producing pathways through synthetic biology, the strains with higher growth rate have a higher potential to be constructed into a microbial cell factory for the overproduction of bulk chemicals. Thus, MG1655 is such a potential strain for the production of bulk chemicals. Furthermore, during the industrial fermentation, high fermentation temperature could lower the cost, the accumulated by-products and toxic compounds might cause the acid and osmotic shocks. Therefore, a strain with high resistant to the heat-, osmotic- and acid-shock will be more robust and economical during the fermentation processes. Our results showed that MG1655 has the stronger tolerance to the heat-, acid-, and osmotic stress than that of other strains, and indicated that some MG1655 derivatives might be engineered into a productive and robust platform for bulk chemicals overproduction.

### 3.2. Construction a MG1655 derivative with increased transformation efficiency

Compared with the seven commercial strains described above, wild-type MG1655 has a lower transformation efficiency, around 10^3^ (cfu/μg DNA) due to the genetic characteristics [41, 42]. In comparison to the seven strains, MG1655 has no artificial genomic modification (Table. 1). To improve the transformation efficiency, a MG1655 derivative was constructed by deleting the genes involved in the defense system.

**Table 1.** Artificial genomic modifications in eight common microbial cell factories.

| Strains | Artificial genomic modifications |  |  |  |
| --- | --- | --- | --- | --- |
| DH5α | *recA1* | *endA1* | *hsdR17(rK<sup>-</sup>, mK<sup>+</sup>)* |  |
| DH10B | *recA1* | *endA1* | $\Delta(mrr\text{-}hsdRMS\text{-}mcrBC)$ | *mcrA* |
| TOP10 | *recA1* | *endA1* | $\Delta(mrr\text{-}hsdRMS\text{-}mcrBC)$ | *mcrA* |
| XL1-blue | *recA1* | *endA1* | *hsdR17(rK<sup>-</sup>, mK<sup>+</sup>)* |  |
| XL10-gold | *recA1* | *endA1* | $\Delta(mcrCB\text{-}hsdSMR\text{-}mrr)173$ | $\Delta(mcrA)183$ |
| BL21(DE3) |  |  | *hsdSB(rB<sup>-</sup>mB<sup>-</sup>)* |  |
| Mach1-T1 | $\Delta recA1398$ | *endA1* | *hsdR (rK<sup>-</sup>, mK<sup>+</sup>)* | |
| MG1655 |  |  | none |  |

The whole process of genome editing included the replacement of the gene cluster *araB-araA-araD* (*araBAD*) with the tetracycline resistance gene (*tet*^r^), deletion of the R-M system of MG1655, and inactivation of genes *mcrA*, *endA*, and *recA* (Fig. 2a). These five genome modifications of MG1655 could facilitate the circular DNA or linear DNA to cross the cell membrane into the cell, improve the exogenous DNA stability, and the anti-contamination ability of MG1655. All genome modifications were completed precisely by using the CRISPR-Cas9-coupled λ-Red recombineering system [43]. In the CRISPR-Cas9 system, gene *cas9* in the plasmid was under the control of arabinose-inducible promoter. To effectively enhance the induction intensity, prolong the induction time, and increase the anti-contamination ability, the arabinose degradation gene cluster *araBAD* was replaced with *tet*^r^ gene. The inactivation of genes *mcrA*, *endA*, and *recA* were completed by inserting a 20-bp fragment into the target site to make a frame shift mutation. The corresponding gel pictures and sequencing results were shown in Fig S3 and Fig S4. And the detailed gene sequences of modifications were shown in supplementary unit 1. As a result, the five rounds of iterative gene editing changed the genotype of MG1655 and increased the transformation efficiency by 168 times to 1.68×10^5^ (cfu/μg DNA). The transformation efficiency of these five intermediate strains were further investigated (Fig. 2c). As shown in the results, compared with the parental strain MG1655, the deletion of *araBAD* and *mcrBC-hsdSMR-mrr* gene cluster did not have a significant influence on the transformation efficiency, but the transformation efficiency of JW080, JW081, and JW128 were significantly improved. Particularly, inactivating *recA* caused an order of magnitude increase on transformation efficiency from 5.7×10^4^ to 1.68×10^5^ (cfu/μg DNA). In conclusion, the five rounds of gene editing synergistically improved the transformation efficiency.

**Fig. 2.**
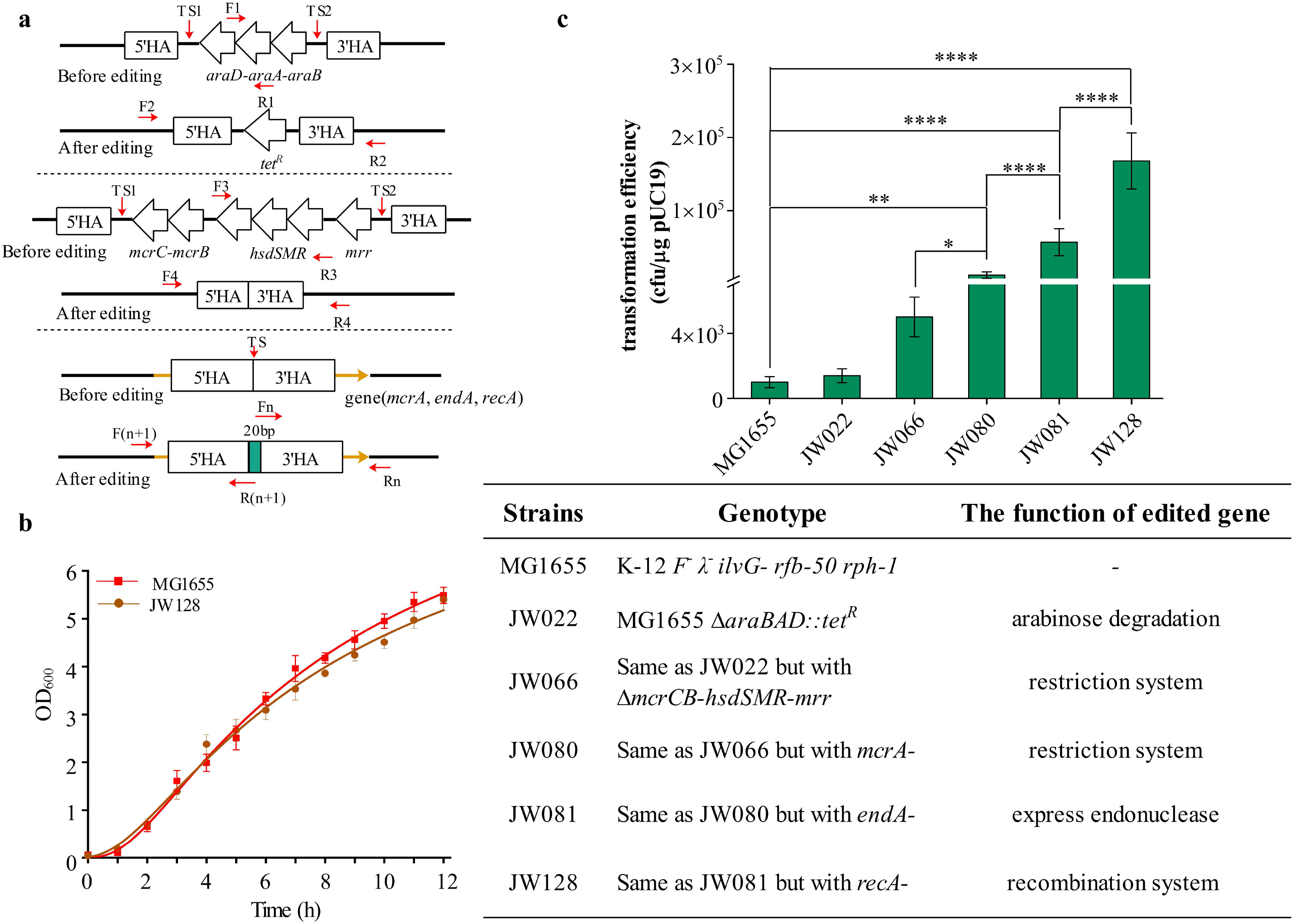
Construction a MG1655 derivative with increased transformation efficiency. (a) The detail gene editing of the modification. (b) The comparison of the growth curve of JW128 and MG1655, the growth curves were fitted by the model of Growth. (c) The transformation efficiency of the strains (from JW022 to JW128) involved in genome editing. The genotypes of corresponding strains were shown in the table. The asterisk on the line represented the significance level between the two columns. More asterisk means greater differences between the two strains. \**P* < 0.1, \*\**P* < 0.01, \*\*\*\**P* < 0.0001 by one‐way ANOVA with Tukey multiple comparison test (P < 0.05). Error bars indicate s.d. (n = 3).

The growth and robustness of engineered strain JW128 were also measured under the unstressed and three stressed conditions as described previously. The fitted growth curve of JW128 was almost the same as for that of MG1655 (Fig. 2b). At 12 h, the OD_600_ of JW128 and MG1655 were 5.4 and 5.49, respectively. According to the growth curves, the specific growth rates of JW128 and MG1655 were 1.2 h^−1^ and 1.4 h^−1^, respectively (Fig. S5). For the robustness, the resistance of JW128 to three stresses decreased a little compared with that of MG1655 because of the bigger relative changes of OD_600_ of JW128, the robustness of JW128 was still maintained, compared with most of the other seven strains (Fig S5). In conclusion, JW128 was an engineering strain with high growth rate, strong robustness, and great anti-contamination ability.

### 3.3. Optimization of the JW128 transformation procedure

Though the transformation efficiency of JW128 had been improved compared with MG1655, it still could not meet the requirements of the existing commercial strains. Thus, the methods were further optimized to make high-efficiency competent cells. The control method was defined as culturing the competent cells in LB medium at 37 °C, 0.1 M CaCl_2_ was used as the wash buffer. Since lower culture temperature [44, 45], metal ions [46, 47], and the hypertonic solution polyethylene glycol (PEG) or DMSO [48–50] could improve the transformation efficiency of *E. coli* competent cells, culture medium (Fig. 3a), culture temperature (Fig. 3b), PEG8000 (40% w/v) (Fig. 3c), and wash buffer (0.1 M CaCl_2_) (Fig. 3d) were adjusted to investigate the optimal methods in our study.

**Fig. 3.**
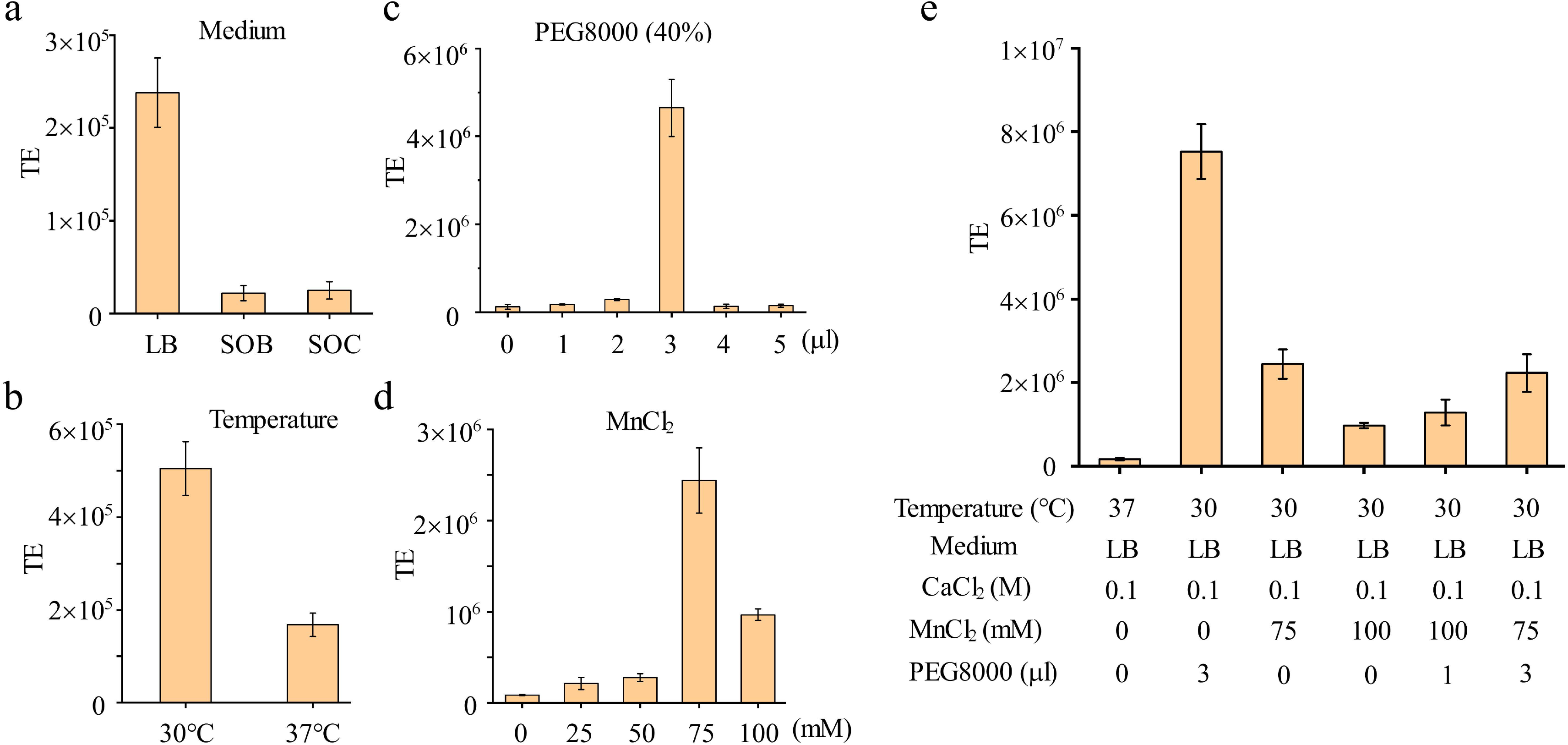
The optimization of methods to make high-efficiency competent cells of JW128. TE: transformation efficiency (cfu/μg DNA). Based on CaCl_2_ method (the first column in e), the transformation efficiency of strain by only changing culture medium (a), culture temperature (b), adding PEG8000 to the mixture of DNA and competent cells when doing the transformation (c), or adding MnCl_2_ to wash buffer (0.1 M CaCl_2_) (d). (e) The transformation efficiency of strains by using optimized methods. Error bars indicate s.d. (n = 3).

Firstly, with three different mediums, the transformation efficiency did not increase with the enrichment of nutrients in the medium (LB, SOB, and SOC medium). LB medium, as the most prevailing one among all the options, was proved to be the optimal medium for the preparation of competent cells (Fig. 3a). Secondly, the transformation efficiency was increased from 1.68×10 ^5^ to 5.05×10 ^5^ (cfu/μg DNA) as the temperature changed from 37 °C to 30 °C (Fig. 3b). When various volumes of PEG8000 was added to the 100 μl mixture of DNA and competent cells, the transformation efficiency kept increasing till the volume of PEG8000 reached 3 μl. The highest efficiency was 4.65×10 ^6^ (cfu/μg DNA), about 38-fold higher than that obtained by the control method (Fig. 3c). Then as the volume of PEG8000 went up to 5 μl, the efficiency decreased to the initial level of the control method. Then the transformation efficiency was measured by using the mixture of CaCl_2_ (0.1 M) and MnCl_2_ with various concentrations (from 25 mM to 100 mM) as wash buffer (Fig. 3d). As shown in the results, the transformation efficiency first increased but then decreased along with the MnCl_2_ concentration increasing to 100 mM. The highest transformation efficiency was 2.44×10 ^6^ (cfu/μg DNA) with 75 mM MnCl_2_ and then the transformation efficiency was decreased to 9.68×10 ^5^ (cfu/μg DNA) with 100 mM MnCl_2_. In conclusion, all experimental groups with the additional MnCl_2_ and CaCl_2_ could improve the transformation efficiency of competent cells compared with the control group. Based on the optimized parameters, the transformation efficiency of competent cells made by five optimized methods were measured (Fig. 3e). The transformation efficiency of all these five methods were increased compared with that of the pre-optimization method. Furthermore, among the five methods, incubating the culture medium at 30 °C and adding 3 microliters PEG8000 (40% w/v) to the mixture of competent cells and DNA fragments was proved to be the optimal method, in which transformation efficiency could reach up to 7.52×10 ^6^ (cfu/μg DNA) and was about 46 times higher than that obtained by the control method and comparable to the commercial strains.

As JW128 possessed the comparable transformation efficiency and the mutations of *endA* and *recA* genes, [13, 51], we speculated that JW128 could also be applied in recombinant plasmids construction. A recombinant plasmid was indeed successfully constructed in JW128 by using T5 exonuclease DNA assembly (TEDA) [52] method in the study (The data not shown). Furthermore, the strains mentioned in the previous reports about making high‐efficiency competent cells mostly were DH5α, XL1-Blue, DH10B, JM109, and BL21 [44, 53, 54], except for MG1655. Our study provided a new sight on improving the transformation efficiency of MG1655.

### 3.4. The higher alcohol production using a JW128 derivative

To verify the potential of JW128 being a microbial cell factory, we introduced the isobutanol synthetic pathway into four strains including MG1655, JW128, JCL16, and JCL260 to produce isobutanol (C4) and methylbutanols (3‐methyl‐1‐butanol and 2‐ methyl‐1‐butanol, C5) (Fig. 4a). JCL16 is the common isobutanol production host strain [55, 56], and JCL260 is currently reported as the best isobutanol producing strain [56, 57].

**Fig. 4.**
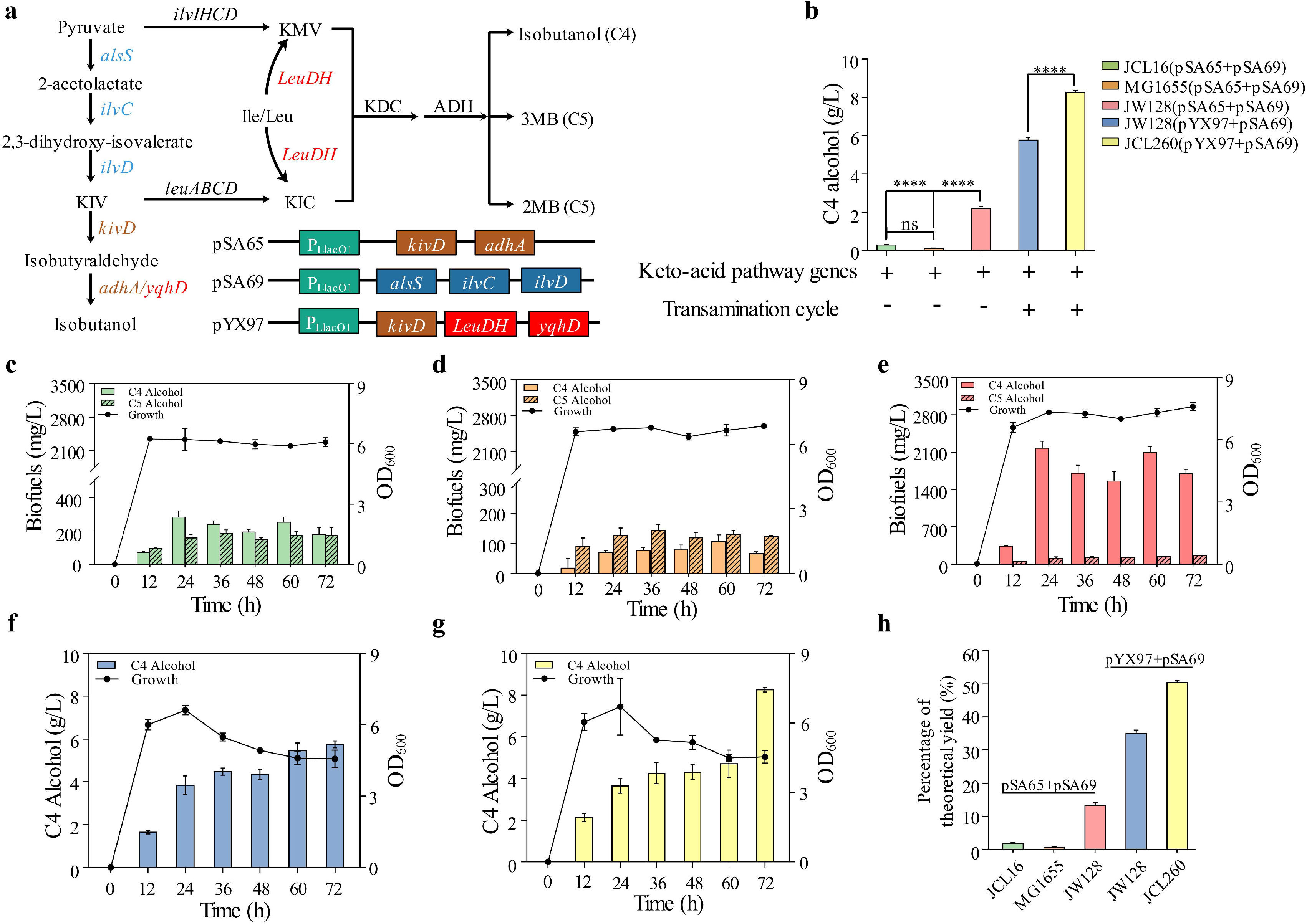
The production of biofuels. 3MB (3‐methyl‐1‐butanol), 2MB (2‐methyl‐1‐ butanol). KIC, 2-ketoisocaproate; KMV, 2-ketomethylvalerate; KDC, 2-keto acid decarboxylase; ADH, alcohol dehydrogenase. (a) Isobutanol (C4) and MB (3MB and 2MB, C5) are produced by synthetic pathway, the essential genes are integrated into three plasmids, they are pSA65, pSA69 and pYX97. (b) The isobutanol production aggregation of strains in panel c to g. \*\*\*\**P* < 0.0001 by one‐way ANOVA with Tukey multiple comparison tests (P < 0.05). (c-e) The biofuels production of strain JCL16, MG1655, JW128 harboring the plasmids pSA69 and pSA65. The isobutanol production of strain JW128 (f) and JCL260 (g) harboring the plasmids pSA69 and pYX97. (h) The percentage of the theoretical yield of isobutanol (C4) produced by five strains. Error bars indicate s.d. (n = 3).

By introducing the keto-acid pathway, the strains JCL16, MG1655, and JW128 harboring plasmids pSA65 and pSA69 were screened for C4 and C5 alcohol production. With 40 g/L glucose, JCL16/pSA65/pSA69 produced 0.283 g/L isobutanol within 24 h (Fig. 4c), yielding a productivity of 0.012 g/L/h and 1.73 % of the theoretical maximum (Fig. 4h). In comparison, MG1655/pSA65/pSA69 produced 0.107 g/L isobutanol at 60 h, which was 2-fold less than JCL16/pSA65/pSA69 (Fig. 4d). Whereas 2.181 g/L isobutanol was produced by JW128/pSA65/pSA69 within 24 h (Fig. 4e), representing 13 % of the theoretical maximum (Fig. 4h). As for the production of C5 alcohol, the titer of JCL16/pSA65/pSA69, MG1655/pSA65/pSA69, and JW128/pSA65/pSA69 were 0.187 g/L, 0.145 g/L, and 0.163 g/L, respectively. From the above data, the C5 alcohol productivity of three strains did not show an significant difference. Opposed to the C5 alcohol production, JW128/pSA65/pSA69 had a nearly 20-fold and 5-fold increase in the productivity of isobutanol production compared with that of MG1655/pSA65/pSA69 and JCL16/pSA65/pSA69, respectively. To explore the isobutanol productivity of JW128 derivative, a transamination and deamination cycle was introduced into the JW128 and JCL260 by overexpressing gene *LeuDH* [55] on plasmid pYX97. Using JCL260/pSA69/pYX97 as the expression host, the titer of isobutanol reached up to 8.26 g/L within 72 h (Fig. 4g) and the productivity was 0.115 g/L/h, which was 50.39 % of the theoretical maximum (Fig. 4h). For the strain JW128/pSA69/pYX97, 5.75 g/L isobutanol was produced within 72 h, reaching a productivity of 0.08 g/L/h and 35.11 % of the theoretical maximum (Fig. 4h).

As a whole (Fig. 4b), isobutanol productivity of MG1655 was subtly weaker (but not significant) than JCL16. However, after the genome editing, the engineering strain JW128 had a significant improvement on isobutanol productivity compared with MG1655 and JCL16. The isobutanol produced by JW128 derivate was almost 5-fold more than that obtained by JCL16 derivate. Compared with JCL260/ pSA69/pYX97, JW128/pSA69/pYX97 had a gap of around 3 g/L in the production of isobutanol. Nevertheless, the deletion of six genes including *adhE*, *ldhA*, *frdBC*, *fnr*, *pta*, and *pflB* operated on JCL260 genome, which highly improved the isobutanol productivity [57–59]. Thus, we showed in this study that JW128 is an alternative host strain for biofuel production.

During the bulk chemicals fermentation, the microbial cell factories usually need to balance the growth maintenance and chemical synthesis and to resist environment stresses. However, once entering the stationary phase or confronting stresses, the cell will tend to allocate more resources to growth maintenance rather than chemical production [60]. Thus, a robust and growth phase-independent host strain urgently required for the value-added chemicals production, redistributing more resources to the producing part. Recently, for the isobutanol production, we discovered the σ^38^-dependent *gadA* promoter [61] and σ^54^-dependent *glnAp2* [60] promoter that could drive the pathway gene overexpression in not only the exponential phase but also in stationary phase, which improved the yield of isobutanol sequentially. Moreover, under the stressed condition, such as low pH and high osmolarity, the pathway enzyme overexpression could still be driven by these two promoters. Therefore, a robust strain (e.g. MG1655) combined with the promoter will be more resistant to the stressed conditions during the fermentation and present a higher productivity in the stationary phase.

### 3.5. The lycopene production using a JW128 derivative

As a type of antioxidant, lycopene plays an important role in reducing the risk of cardiovascular disease, genetic damage, and inhibiting tumor development [62]. In the pre-experiment of lycopene production with seven *E. coli* strains described above, DH5α and DH10B showed higher productivity than that of other strains. Thus, four strains including DH5α, DH10B, MG1655, and JW128 harboring plasmid p‐P_fumAp_‐ *crtEBI* were cultured in LB medium for 22 hours to investigate the productivities of lycopene. Since lycopene would not be secreted outside the cell, whether lycopene had been produced could be directly distinguished from the cells’ colors (Fig. 5a). Among four strains, it can be preliminarily determined that MG1655 is the worst host strain of lycopene production because the color of MG1655/p‐P_fumAp_‐*crtEBI* cells had no observed changes while that of other strains became clearly red (Fig. 5b). The cell biomass of MG1655/p‐P_fumAp_‐*crtEBI* and JW128/p‐P_fumAp_‐*crtEBI* reached the highest level with the OD_600_ reaching up to 4.05 and 4.22 at 22 h respectively (Fig. 5c). The OD_474_ of lycopene extract was measured to calculate the titer of lycopene. The OD_474_ of JW128/p‐P_fumAp_‐*crtEBI* and MG1655/p‐P_fumAp_‐*crtEBI* were 0.746 and 0.154, respectively, which were two extremes of the four strains. Through the conversion with a standard curve, the titer of lycopene produced by MG1655, JW128, DH5α, and DH10B harboring the p‐P_fumAp_‐*crtEBI* plasmid was 0.37 ± 0.07 g/L, 1.91 ± 0.15 g/L, 1.36 ± 0.12 g/L and 1.32 ± 0.07 g/L, respectively (Fig. 5d). As a whole, though the cell biomass of MG1655/p‐P_fumAp_‐*crtEBI* and JW128/p‐P_fumAp_‐*crtEBI* were similar, the titer of lycopene produced by JW128/p‐P_fumAp_‐*crtEBI* was the highest and almost 6 times of that obtained in MG1655/p‐P_fumAp_‐*crtEBI*. It can be speculated that the plasmid in JW128 expressed much stronger than in MG1655.

**Fig. 5.**
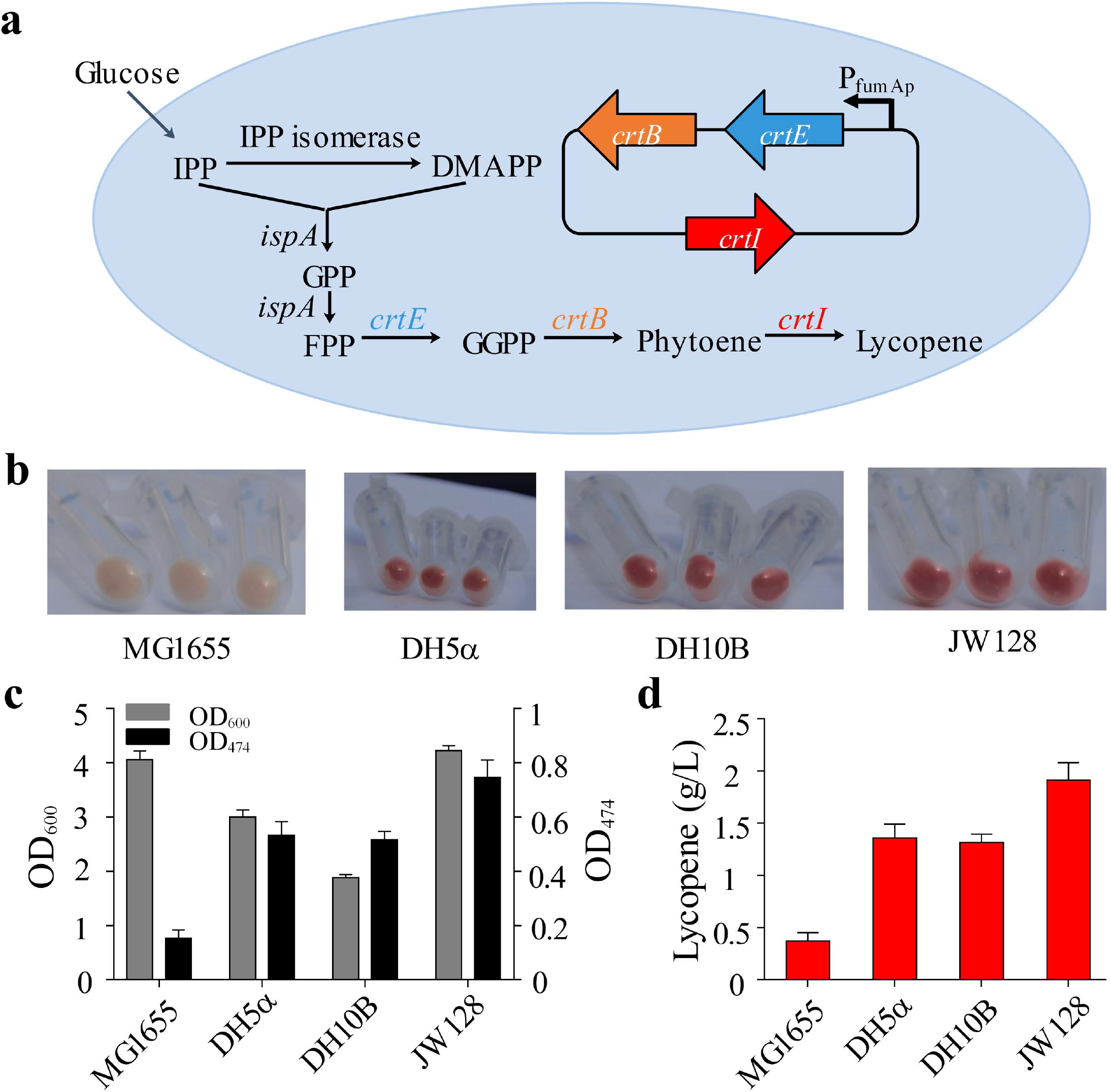
The lycopene production of different strains in LB medium. (a) The production pathway of lycopene in *E. coli*, the genes in the ellipse are provided by the plasmid used in this study. IPP (Isopentyl pyrophosphate), DMAPP (dimethylallyl pyrophosphate), GPP (Geranyl pyrophosphate), FPP (farnesyl pyrophosphate), GGPP (Geranylgeranyl pyrophosphate). (b) The cell pellet of 4 mL bacteria solution of four strains after cultured for 22 hours. (c) The value of OD_600_ and OD_474_ of four strains. OD_600_ is measured to show the cell biomass, OD_474_ is used to qualify the lycopene content. (d) The lycopene yield of four strains, which converted by the standard curve. Error bars indicate s.d. (n = 3).

In the previous study, to realize the efficient chemical production in MG1655, modification of the corresponding producing pathway to compensate for the low expression of plasmids was more labor-consuming (Table. 2), In our study, we turned to the modification of MG1655 genome to improve the plasmid stability and producing capacity, which engineered the MG1655 to be a universal chassis host. Thus, with this strategy and the promoters described above, MG1655 derivatives might be transformed into an optimal platform for value-added chemicals overproduction.

**Table 2.** The chemicals produced by MG1655 or MG1655 derivatives.

|  | Chemicals | Titer (g/L) | Approach | reference |
| --- | --- | --- | --- | --- |
| Acid | Succinic acid | 42.5 | Deletion of *pflB* and *ldhA*. Overexpression of PEP carboxykinase. | [63] |
|  | Itaconic acid | 47 | Using temperature-sensitive promoter to control the TCA cycle dynamically | [64] |
|  | Citramalic acid | 46.5 | Deletion of citrate synthase and acetate kinase | [65] |
|  | Fatty acid | 82.64 (mg/g) | Overexpression of *accA*, *accB*, *accC*, *fabD*, and EC 3.1.2.14 gene | [66] |
| Amino acid | L-serine | 11.7 | Deletion of three L-serine deaminases (gene *sdaA*, *sdaB*, and *tdcG*) and serine hydroxyl methyl transferase (gene *glyA*). Overexpression of *eamA* | [67] |
| Alcohol | 1,4-butanediol | 110 | Containing Cat2 variants with improved BDO tolerance and Ald variant with improved *in vitro* activity, expression, and stability | [68] |
|  | 1,2-Propanediol | 5.6 | Overexpression of *mgsA*, *gldA*, and *yqhD*. Replacing the native PEP-dependent dihydroxyacetone kinase (DHAK) with an ATP-dependent DHAK. Disrupting the lactate and acetate synthesis pathway. | [69] |
|  | Ethylene glycol | 20 | Deletion of *xylB*, *aldA*. Overexpressing *fucO* | [70] |
| Terpenoid | Taxadiene | 1 | Separating the taxadiene metabolic pathway into two modules | [71] |
| Aldehyde | Vanillin | 0.119 | Deletion of native aldehyde reductases and expression of key enzymes for converting 3-dehydroshikimate to vanillin | [72] |
| Ester | 2-phenylethyl<br>acetate | 0.268 | Combined the synthesis pathway of 2-phenylethanol and 2-phenylethylacetate by overexpressing an alcohol acetyltransferase ATF1 | [73] |

## 4. Conclusions

In this work, an engineering strain JW128 was constructed by replacing *araBAD* with *tet*^r^, deleting *mcrBC-hsdSMR-mrr*, and inactivating *mcrA*, *endA*, and *recA*. JW128 not only kept the growth and robustness advantage of MG1655 but also equipped with the enhanced transformation efficiency and plasmid stability. JW128, as a robust and universal chassis host, had the potential to be applied to construct recombinant plasmid and produce the desired chemicals such as isobutanol and lycopene.

## Ethics approval and consent to participate

Not applicable.

## Consent for publication

Not applicable.

## Availability of data and materials

All the data generated during the current study are included in the manuscript.

## Author contributions

Jingge Wang: Conceptualization, Methodology, Formal analysis, Investigation, Writing-Original Draft. Chaoyong Huang: Conceptualization, Methodology, Formal analysis. Kai Guo: Methodology. Lianjie Ma: Investigation. Xiangyu Meng: Investigation. Ning Wang: Writing-Original Draft. Yi-Xin Huo: Conceptualization, Methodology, Formal analysis, Writing-Original Draft.

## Acknowledgments

The work finished in Beijing Institute of Technology was supported by the National Natural Science Foundation of China (Grant No. 21676026), the National Key R&D Program of China (grant No. 2017YFD0201400), the Fundamental Research Funds for the Central Universities.

## References

1. Fang H, Li D, Kang J, Jiang P, Sun J, Zhang D: Metabolic engineering of *Escherichia coli* for de novo biosynthesis of vitamin B12. Nat Commun 2018; 9:4917. doi:10.1038/s41467-018-07412-6

2. Yu Y, Zhu X, Xu H, Zhang X: Construction of an energy-conserving glycerol utilization pathways for improving anaerobic succinate production in *Escherichia coli*. Metab Eng 2019; 56:181–189. doi:10.1016/j.ymben.2019.10.002

3. Park SY, Binkley RM, Kim WJ, Lee MH, Lee SY: Metabolic engineering of *Escherichia coli* for high-level astaxanthin production with high productivity. Metab Eng 2018; 49:105–115. doi:10.1016/j.ymben.2018.08.002

4. Monk JM, Koza A, Campodonico MA, Machado D, Seoane JM, Palsson BO, Herrgard MJ, Feist AM: Multi-omics Quantification of Species Variation of *Escherichia coli* Links Molecular Features with Strain Phenotypes. Cell systems 2016; 3:238–251.

5. Archer CT, Kim JF, Jeong H, Jin HP, Nielsen LK: The genome sequence of *E. coli* W (ATCC 9637): Comparative genome analysis and an improved genome-scale reconstruction of *E. coli*. Bmc Genomics 2011; 12:9.

6. Arifin Y, Archer C, Lim S, Quek LE, Sugiarto H, Marcellin E, Vickers CE, Kromer JO, Nielsen LK: *Escherichia coli* W shows fast, highly oxidative sucrose metabolism and low acetate formation. Appl Microbiol Biotechnol 2014; 98:9033–9044. doi:10.1007/s00253-014-5956-4

7. Yoon SH, Han M-J, Jeong H, Lee CH, Xia X-X, Lee D-H, Shim JH, Lee SY, Oh TK, Kim JF: Comparative multi-omics systems analysis of *Escherichia coli* strains B and K-12. Genome Biology; 13:R37.

8. Vijayendran C, Polen T, Wendisch VF, Friehs K, Niehaus K, Flaschel E: The plasticity of global proteome and genome expression analyzed in closely related W3110 and MG1655 strains of a well-studied model organism, *Escherichia coli*-K12. J Biotechnol 2007; 128:747–761. doi:10.1016/j.jbiotec.2006.12.026

9. Chae HS, Kim KH, Kim SC, Lee PC: Strain-dependent carotenoid productions in metabolically engineered *Escherichia coli*. Appl Biochem Biotechnol 2010; 162:2333–2344. doi:10.1007/s12010-010-9006-0

10. Marisch K, Bayer K, Scharl T, Mairhofer J, Krempl PM, Hummel K, Razzazi-Fazeli E, Striedner G: A comparative analysis of industrial *Escherichia coli* K-12 and B strains in high-glucose batch cultivations on process-, transcriptome- and proteome level. PLoS One 2013; 8:e70516. doi:10.1371/journal.pone.0070516

11. Blattner FR, Plunkett G, ., Bloch CA, Perna NT, Burland V, ., Riley M, ., Collado-Vides J, ., Glasner JD, Rode CK, Mayhew GF: The complete genome sequence of *Escherichia coli* K-12. Science 1997; 277:1453–1462.

12. Durfee T, Nelson R, Baldwin S, Plunkett G, 3rd, Burland V, Mau B, Petrosino JF, Qin X, Muzny DM, Ayele M, et al: The complete genome sequence of *Escherichia coli* DH10B: insights into the biology of a laboratory workhorse. J Bacteriol 2008; 190:2597–2606. doi:10.1128/jb.01695-07

13. Phue JN, Lee SJ, Trinh L, Shiloach J: Modified *Escherichia coli* B (BL21), a superior producer of plasmid DNA compared with *Escherichia coli* K (DH5alpha). Biotechnol Bioeng 2008; 101:831–836. doi:10.1002/bit.21973

14. Loenen WA: Tracking EcoKI and DNA fifty years on: a golden story full of surprises. Nucleic Acids Res 2003; 31:7059–7069. doi:10.1093/nar/gkg944

15. Murray NE: Type I restriction systems: sophisticated molecular machines (a legacy of Bertani and Weigle). Microbiol Mol Biol Rev 2000; 64:412–434. doi:10.1128/mmbr.64.2.412-434.2000

16. Sain B, Murray NE: The hsd (host specificity) genes of *E. coli* K 12. Mol Gen Genet 1980; 180:35–46. doi:10.1007/bf00267350

17. Raleigh EA, Wilson G: *Escherichia coli* K-12 restricts DNA containing 5-methylcytosine. Proc Natl Acad Sci U S A 1986; 83:9070–9074. doi:10.1073/pnas.83.23.9070

18. Fleischman RA, Cambell JL, Richardson CC: Modification and restriction of T-even bacteriophages. In vitro degradation of deoxyribonucleic acid containing 5-hydroxymethylctosine. J Biol Chem 1976; 251:1561–1570.

19. Ross TK, Achberger EC, Braymer HD: Identification of a second polypeptide required for McrB restriction of 5-methylcytosine-containing DNA in *Escherichia coli* K12. Mol Gen Genet 1989; 216:402–407. doi:10.1007/bf00334382

20. Dila D, Sutherland E, Moran L, Slatko B, Raleigh EA: Genetic and sequence organization of the *mcrBC* locus of *Escherichia coli* K-12. J Bacteriol 1990; 172:4888–4900. doi:10.1128/jb.172.9.4888-4900.1990

21. Waite-Rees PA, Keating CJ, Moran LS, Slatko BE, Hornstra LJ, Benner JS: Characterization and expression of the Escherichia coli Mrr restriction system. J Bacteriol 1991; 173:5207–5219. doi:10.1128/jb.173.16.5207-5219.1991

22. Sharan SK, Thomason LC, Kuznetsov SG, Court DL: Recombineering: a homologous recombination-based method of genetic engineering. Nat Protoc 2009; 4:206–223. doi:10.1038/nprot.2008.227

23. Jeong J, Cho N, Jung D, Bang D: Genome-scale genetic engineering in *Escherichia coli*. Biotechnol Adv 2013; 31:804–810. doi:10.1016/j.biotechadv.2013.04.003

24. Pines G, Freed EF, Winkler JD, Gill RT: Bacterial Recombineering: Genome Engineering via Phage-Based Homologous Recombination. ACS Synth Biol 2015; 4:1176–1185. doi:10.1021/acssynbio.5b00009

25. Wang HH, Isaacs FJ, Carr PA, Sun ZZ, Xu G, Forest CR, Church GM: Programming cells by multiplex genome engineering and accelerated evolution. Nature 2009; 460:894–898. doi:10.1038/nature08187

26. Warner JR, Reeder PJ, Karimpour-Fard A, Woodruff LB, Gill RT: Rapid profiling of a microbial genome using mixtures of barcoded oligonucleotides. Nat Biotechnol 2010; 28:856–862. doi:10.1038/nbt.1653

27. Isaacs FJ, Carr PA, Wang HH, Lajoie MJ, Sterling B, Kraal L, Tolonen AC, Gianoulis TA, Goodman DB, Reppas NB, et al: Precise manipulation of chromosomes in vivo enables genome-wide codon replacement. Science 2011; 333:348–353. doi:10.1126/science.1205822

28. Datsenko KA, Wanner BL: One-step inactivation of chromosomal genes in *Escherichia coli* K-12 using PCR products. Proc Natl Acad Sci U S A 2000; 97:6640–6645. doi:10.1073/pnas.120163297

29. Tischer BK, von Einem J, Kaufer B, Osterrieder N: Two-step red-mediated recombination for versatile high-efficiency markerless DNA manipulation in *Escherichia coli*. Biotechniques 2006; 40:191–197. doi:10.2144/000112096

30. Yu BJ, Kang KH, Lee JH, Sung BH, Kim MS, Kim SC: Rapid and efficient construction of markerless deletions in the *Escherichia coli* genome. Nucleic Acids Res 2008; 36:e84. doi:10.1093/nar/gkn359

31. Yang J, Sun B, Huang H, Jiang Y, Diao L, Chen B, Xu C, Wang X, Liu J, Jiang W, Yang S: High-efficiency scarless genetic modification in *Escherichia coli* by using lambda red recombination and I-SceI cleavage. Appl Environ Microbiol 2014; 80:3826–3834. doi:10.1128/aem.00313-14

32. Urnov FD, Rebar EJ, Holmes MC, Zhang HS, Gregory PD: Genome editing with engineered zinc finger nucleases. Nat Rev Genet 2010; 11:636–646. doi:10.1038/nrg2842

33. Joung JK, Sander JD: TALENs: a widely applicable technology for targeted genome editing. Nat Rev Mol Cell Biol 2013; 14:49–55. doi:10.1038/nrm3486

34. Bogdanove AJ, Voytas DF: TAL effectors: customizable proteins for DNA targeting. Science 2011; 333:1843–1846. doi:10.1126/science.1204094

35. Wood AJ, Lo TW, Zeitler B, Pickle CS, Ralston EJ, Lee AH, Amora R, Miller JC, Leung E, Meng X, et al: Targeted genome editing across species using ZFNs and TALENs. Science 2011; 333:307. doi:10.1126/science.1207773

36. Jinek M, Chylinski K, Fonfara I, Hauer M, Doudna JA, Charpentier E: A programmable dual-RNA-guided DNA endonuclease in adaptive bacterial immunity. Science 2012; 337:816–821. doi:10.1126/science.1225829

37. Gasiunas GB, R. Horvath, P. Siksnys, V.: Cas9-crRNA ribonucleoprotein complex mediates specific DNA cleavage for adaptive immunity in bacteria. Proc Natl Acad Sci U S A 2012; 109:E2579–2586. doi:10.1073/pnas.1208507109

38. Jiang W, Bikard D, Cox D, Zhang F, Marraffini LA: RNA-guided editing of bacterial genomes using CRISPR-Cas systems. Nat Biotechnol 2013; 31:233–239. doi:10.1038/nbt.2508

39. Yu H, Wang N, Huo W, Zhang Y, Zhang W, Yang Y, Chen Z, Huo YX: Establishment of BmoR-based biosensor to screen isobutanol overproducer. Microb Cell Fact 2019; 18:30.

40. Kim SW, Keasling JD: Metabolic engineering of the nonmevalonate isopentenyl diphosphate synthesis pathway in *Escherichia coli* enhances lycopene production. Biotechnol Bioeng 2001; 72:408–415. doi:10.1002/1097-0290(20000220)72:4<408::aid-bit1003>3.0.co;2-h

41. Sambrook J, Russell DW: The inoue method for preparation and transformation of competent *E. Coli*: “ultra-competent” cells. Csh Protoc 2006; 2006:418–424.

42. Arunasri K, Adil M, Charan KV, Suvro C, Reddy SH, Shivaji S: Effect of simulated microgravity on *E. coli* K12 MG1655 growth and gene expression. PLOS ONE 2013; 8.

43. Jiang Y, Chen B, Duan C, Sun B, Yang J, Yang S: Multigene editing in the *Escherichia coli* genome via the CRISPR-Cas9 system. Appl Environ Microbiol 2015; 81:2506–2514. doi:10.1128/aem.04023-14

44. Inoue H, Nojima H, Okayama H: High efficiency transformation of *Escherichia coli* with plasmids. Gene 1990; 96:23–28.

45. Hanahan D, Jessee J, Bloom FR: Plasmid transformation of *Escherichia coli* and other bacteria. Methods Enzymol 1991; 204:63–113. doi:10.1016/0076-6879(91)04006-a

46. Aune TE, Aachmann FL: Methodologies to increase the transformation efficiencies and the range of bacteria that can be transformed. Appl Microbiol Biotechnol 2010; 85:1301–1313. doi:10.1007/s00253-009-2349-1

47. Hanahan D: Studies on transformation of *Escherichia coli* with plasmids. J Mol Biol 1983; 166:557–580. doi:10.1016/s0022-2836(83)80284-8

48. Kurien BT, Scofield RH: Polyethylene glycol-mediated bacterial colony transformation. BioTechniques 1995; 18:1023–1026.

49. Chung CT, Miller RH: A rapid and convenient method for the preparation and storage of competent bacterial cells. Nucleic Acids Res 1988; 16:3580. doi:10.1093/nar/16.8.3580

50. Klebe RJ, Harriss JV, Hanson DP, Gauntt CJ: High-efficiency polyethylene glycol-mediated transformation of mammalian cells. Somat Cell Mol Genet 1984; 10:495–502. doi:10.1007/bf01534854

51. Taylor RG, Walker DC, McInnes RR: *E. coli* host strains significantly affect the quality of small scale plasmid DNA preparations used for sequencing. Nucleic Acids Res 1993; 21:1677–1678. doi:10.1093/nar/21.7.1677

52. Xia Y, Li K, Li J, Wang T, Gu L, Xun L: T5 exonuclease-dependent assembly offers a low-cost method for efficient cloning and site-directed mutagenesis. Nucleic Acids Res 2019; 47:e15. doi:10.1093/nar/gky1169

53. Chan WT, Verma CS, Lane DP, Gan SK: A comparison and optimization of methods and factors affecting the transformation of *Escherichia coli*. Biosci Rep 2013; 33. doi:10.1042/bsr20130098

54. Tu Z, He G, Li KX, Chen M, Chang J, Chen L, Yao Q, Liu DP, Ye H, Shi J: An improved system for competent cell preparation and high efficiency plasmid transformation using different *Escherichia coli* strains. Electron J Biotechnol 2005; 8:113–120.

55. Huo Y, Cho KM, Rivera JGL, Monte E, Shen CR, Yan Y, Liao JC: Conversion of proteins into biofuels by engineering nitrogen flux. Nat Biotechnol 2011; 29:346–351.

56. Baez A, Cho KM, Liao JC: High-flux isobutanol production using engineered *Escherichia coli*: a bioreactor study with in situ product removal. Appl Microbiol Biotechnol 2011; 90:1681–1690.

57. Baez A, Cho KM, Liao JC: High-flux isobutanol production using engineered Escherichia coli: a bioreactor study with in situ product removal. Appl Microbiol Biotechnol 2011; 90:1681–1690. doi:10.1007/s00253-011-3173-y

58. Atsumi S, Cann AF, Connor MR, Shen CR, Smith KM, Brynildsen MP, Chou KJ, Hanai T, Liao JC: Metabolic engineering of Escherichia coli for 1-butanol production. Metab Eng 2008; 10:305–311. doi:10.1016/j.ymben.2007.08.003

59. Atsumi S, Hanai T, Liao JC: Non-fermentative pathways for synthesis of branched-chain higher alcohols as biofuels. Nature 2008; 451:86–89. doi:10.1038/nature06450

60. Ma L, Guo L, Yang Y, Guo K, Yan Y, Ma X, Huo YX: Protein-based biorefining driven by nitrogen-responsive transcriptional machinery. Biotechnol Biofuels 2020; 13:29. doi:10.1186/s13068-020-1667-5

61. Huo YX, Guo L, Ma X: Biofuel production with a stress-resistant and growth phase-independent promoter: mechanism revealed by in vitro transcription assays. Appl Microbiol Biotechnol 2018; 102:2929–2940. doi:10.1007/s00253-018-8809-8

62. Chen D, Huang C, Chen Z: A review for the pharmacological effect of lycopene in central nervous system disorders. Biomed Pharmacother 2019; 111:791–801. doi:10.1016/j.biopha.2018.12.151

63. Li Q, Huang B, Wu H, Li Z, Ye Q: Efficient anaerobic production of succinate from glycerol in engineered *Escherichia coli* by using dual carbon sources and limiting oxygen supply in preceding aerobic culture. Bioresour Technol 2017; 231:75–84. doi:10.1016/j.biortech.2017.01.051

64. Harder BJ, Bettenbrock K, Klamt S: Temperature-dependent dynamic control of the TCA cycle increases volumetric productivity of itaconic acid production by *Escherichia coli*. Biotechnol Bioeng 2018; 115:156–164. doi:10.1002/bit.26446

65. Wu X, Eiteman MA: Production of citramalate by metabolically engineered *Escherichia coli*. Biotechnol Bioeng 2016; 113:2670–2675. doi:10.1002/bit.26035

66. Jeon E, Lee S, Won JI, Han SO, Kim J, Lee J: Development of *Escherichia coli* MG1655 strains to produce long chain fatty acids by engineering fatty acid synthesis (FAS) metabolism. Enzyme Microb Technol 2011; 49:44–51. doi:10.1016/j.enzmictec.2011.04.001

67. Mundhada H, Schneider K, Christensen HB, Nielsen AT: Engineering of high yield production of L-serine in *Escherichia coli*. Biotechnol Bioeng 2016; 113:807–816. doi:10.1002/bit.25844

68. Burgard A, Burk MJ, Osterhout R, Van Dien S, Yim H: Development of a commercial scale process for production of 1,4-butanediol from sugar. Curr Opin Biotechnol 2016; 42:118–125. doi:10.1016/j.copbio.2016.04.016

69. Clomburg JM, Gonzalez R: Metabolic engineering of *Escherichia coli* for the production of 1,2-propanediol from glycerol. Biotechnol Bioeng 2011; 108:867–879. doi:10.1002/bit.22993

70. Alkim C, Cam Y, Trichez D, Auriol C, Spina L, Vax A, Bartolo F, Besse P, Francois JM, Walther T: Optimization of ethylene glycol production from (D)-xylose via a synthetic pathway implemented in *Escherichia coli*. Microb Cell Fact 2015; 14:127. doi:10.1186/s12934-015-0312-7

71. Ajikumar PK, Xiao WH, Tyo KE, Wang Y, Simeon F, Leonard E, Mucha O, Phon TH, Pfeifer B, Stephanopoulos G: Isoprenoid pathway optimization for Taxol precursor overproduction in *Escherichia coli*. Science 2010; 330:70–74. doi:10.1126/science.1191652

72. Kunjapur AM, Tarasova Y, Prather KL: Synthesis and accumulation of aromatic aldehydes in an engineered strain of *Escherichia coli*. J Am Chem Soc 2014; 136:11644–11654. doi:10.1021/ja506664a

73. Guo D, Zhang L, Pan H, Li X: Metabolic engineering of *Escherichia coli* for production of 2-Phenylethylacetate from L-phenylalanine. Microbiologyopen 2017; 6. doi:10.1002/mbo3.486

