## supplementary File for "Converting *Escherichia coli* MG1655 into a chemical overproducer through inactivating defense system against exogenous DNA"

### Supplementary information

**Table S1. The reagents and mediums used in this study**

| **Reagent** | **Stock concentration** | **Working concentration** |
| --- | --- | --- |
| Ampicillin | 100 mg/mL | 100 μg/mL |
| Kanamycin | 50 mg/mL | 50 μg/mL |
| Glucose | 500 mg/mL | 10 mg/mL |
| L-arabinose | 1 M | 20 mM |
| IPTG | 100 mM | 0.1 mM |
| **Mediums** | **Components** | |
| Luria-Bertani (LB) | 10 g/L tryptone, 5 g/L yeast extract, 10 g/L NaCl | |
| SOC | 20 g/L tryptone, 5 g/L yeast extract,0.5 g/L NaCl,  0.25 mM KCl, 10 mM MgCl2, 20 mM glucose | |
| SOB | 20 g/L tryptone, 5 g/L yeast extract, 0.5 g/L NaCl,  0.25 mM KCl, 10 mM MgCl2 | |
| M9Y | 11.28 g/L M9 minimal salt, 1 mM MgSO4, 0.1 mM CaCl2,  10 mg/L VB1, 40 g/L glucose and 5 g/L Yeast exact | |

1

### Table S2. The OD600 of eight strains cultured in different conditions

| **Max OD** | | | | |
| --- | --- | --- | --- | --- |
| **Strains** |  |  | **Stressed conditions** | |
| **Unstressed condition** | | | | |
|  |  | **42 ℃** | **pH 5.5** | **Salinity (30 g/L NaCl)** |
| MG1655 | 5.49 | 5.63 | 5.46 | 4.62 |
| DH10B | 2.99 | 2.61 | 2.35 | 0.23 |
| XL10-gold | 3.78 | 3.07 | 3.51 | 1.25 |
| DH5α | 3.64 | 3.83 | 3.73 | 2.21 |
| TOP10 | 3.20 | 2.62 | 3.02 | 2.34 |
| Mach1-T1 | 3.85 | 2.81 | 3.20 | 2.23 |
| XL1-blue | 4.64 | 3.59 | 4.10 | 2.57 |
| BL21(DE3) | 5.01 | 4.12 | 4.79 | 3.01 |

**Table S3. The plasmids used in this study.**

| Plasmids | Descriptions | Sources |
| --- | --- | --- |
| p-PBAD-sgRNA-*recA* | pSC101 ori; *amp^r^*; inactivate *recA* | This study |
| p-PBAD-sgRNA-*endA* | pSC101 ori; *amp^r^*; inactivate *endA* | This study |
| p-PBAD-sgRNA-*mcrA* | pSC101 ori; *amp^r^*; inactivate *mcrA* | This study |
| p-PBAD-sgRNA-*mcrCB-hsdSMR-mrr* | pSC101 ori; *amp^r^*; knock out *mcrCB-hsdSMR-mrr* gene cluster | This study |
| p-PBAD-sgRNA-*araD-araA-araB* | pSC101 ori; *amp^r^*; replace *araD-araA-araB* with *tet^r^* | This study |
| p-PBAD-*cas9*-PT5-Redγαβ | Recombinant plasmid; c*as9;* λ-Red system; p15A ori; *kan^r^* | This study |
| pSA69 | *P*LlacO1-*alsS*-*ilvC*-*ilvD*; p15A; *kan^r^* | [1] |
| pSA65 | *P*LlacO1-*kivd*-*adhA*; p15A; *amp^r^* | [1] |
| pYX97 | *P*LlacO1-*LeuDH*-*alsS*-*ilvC*-*ilvD*; p15A; *kan^r^* | [1] |
| p-PfumAp-crtEBI | pBR322 ori; *amp^r^*; produce lycopene | This study |

### Table S4. The strains used in this study.

| Strains | Description | Reference or Source |
| --- | --- | --- |
| DH5α | F^–^ Φ80*lacZ*Δ*M*15 Δ(*lacZYA-argF*)U169 *recA*1 *endA*1*hsdR*17 (rK–, mKþ) *phoA supE*44 λ  – *thi*-1 *gyrA*96 *relA*1 | Laboratory stock |
| MG1655 | K-12; F^-^λ^-^*rph*-1 | Laboratory stock |
| DH10B | F^–^ *endA1 deoR ^+^ recA1 galE15 galK16 nupG rpsL Δ(lac)X74 φ80lacZΔM15 araD139 Δ(ara,leu)7697 mcrA Δ(mrr-hsdRMS-mcrBC) StrR λ^–^* | Laboratory stock |
| Mach1-T1 | str. W Δ*recA1398 endA1 fhuA* Φ80Δ(*lac*)M15 Δ*(lac)X74 hsdR* (*rK^–^mK^+^*) | Laboratory stock |
| XL10-gold | *endA*1 *glnV*44 *recA*1 *thi*-1 *gyrA*96 *relA*1 lac Hte Δ(*mcrA*)183 Δ(*mcrCB-hsdSMR-mrr*)173  *tet^R^* F'[*proAB lacI^q^ZΔM15 Tn10(Tet^R^ Amy Cm^R^*)] | Laboratory stock |

| **Table S4 (continued)** | | |
| --- | --- | --- |
| Strains | Description | Reference or Source |
| BL21(DE3) | *E. coli* str. B F – *ompT gal dcm lon hsdSB*(*rB*^–^*mB*^–^) λ (DE3 [*lacI lacUV5*-*T7p07 ind1 sam7*  *nin5*]) [*malB*^+^] K-12(λ^S^) | Laboratory stock |
| TOP10 | F^-^ *mcrA* Δ(*mrr-hsdRMS-mcrBC*) φ80*lacZ*ΔM15 Δ*lacX*74 *nupG recA*1 *araD*139 Δ(*ara-*  *leu*)7697 *galE*15 *galK*16 *rpsL*(Str^R^) *endA*1 λ^-^ | Laboratory stock |
| XL1-blue | *endA*1 *gyrA*96(*nal ^R^*) *thi*-1 *recA*1 *relA*1 *lac glnV*44 F'[::Tn10 *proAB*^+^ *lacI*^q^ Δ(*lacZ*)M15]  *hsdR*17(*rK^-^ mK^+^*) | Laboratory stock |
| JW080 | MG1655Δ *araBAD*::*tet^R^* | This study |
| JW081 | MG1655Δ*araBAD*::*tet^R^,* Δ*mcrCB-hsdSMR-mrr* | This study |

| **Table S4 (continued)** | | |
| --- | --- | --- |
| Strains | Description | Reference or Source |
| JW066 | MG1655Δ*araBAD*::*tet^R^,* Δ*mcrCB‐hsdSMR-mrr, mcrA*- | This study |
| JW022 | MG1655Δ*araBAD::tet^R^,* Δ*mcrCB-hsdSMR-mrr, mcrA*-, *endA-* | This study |
| JW128 | MG1655Δ*araBAD*::*tet^R^,* Δ*mcrCB-hsdSMR-mrr, mcrA*-, *endA-, recA-* | This study |
| JCL16 | BW25113/F'[traD36, *proAB*+, *lacIq Z*Δ*M*15] | [2] |
| JCL260 | Same as JCL16 but with Δ*adhE,* Δ*ldhA,* Δ*frdBC,* Δ*fnr,* Δ*pta,* Δ*pflB* | [2] |

**Table S5. The main primers used in this study**

| **Primers** | **Sequence** | **Characteristics** |
| --- | --- | --- |
| F1 R1 F2  R2 | gtcgcatcaggcgttacata gatcgccgcagaagagaaac tcaatctgcccggcaaaca  ctggcgggaacagcaaaata | Colony PCR identification of *araB-araA-araD*  editing |
| F3 R3 F4  R4 | ggtggaataacactcttgcc agcaacaacgttcgcaatcc ccgatcgctatctgcaaaac  caacaccaacggttttatgc | Colony PCR identification of *mcrCB-hsdSMR-*  *mrr* editing |
| F5 R5 F6  R6 | agacaagataagcggcacaa CGACTCTCGATAGAACCACT AGTGGTTCTATCGAGAGTCG  cacctgaagacaggggaatt | Colony PCR identification of *mcrA* editing |
| F7 R7 F8  R8 | ctatcctctgccgctatgaa GCAACACAGAAGGGGTACAT ATGTACCCCTTCTGTGTTGC  cgttgcacatacgggttatg | Colony PCR identification of *endA* editing |
| F9 R9 F10  R10 | gcgtagaatttcagcgcgtta gGGCGATACTATGAGAAGACC gGGTCTTCTCATAGTATCGCC  ctgacagtgaactgatgcag | Colony PCR identification of *recA* editing |
| iBT_Huolab-468hcy-F1 iBT_Huolab-509hcy-R1 iBT_Huolab-510hcy-F2 iBT_Huolab-511hcy-R2 iBT_Huolab-629hcy-F3  iBT_Huolab-1056hcy-R3 | agataggtgcctcactgatt cggaaggatctgaggttctt gtctgctatgtggtgctatct tgccacctgacgtctaagaa cgagccggaagcataaagt  cgcgtaccatgggatcctta | Colony PCR identification of B series plasmid construction |

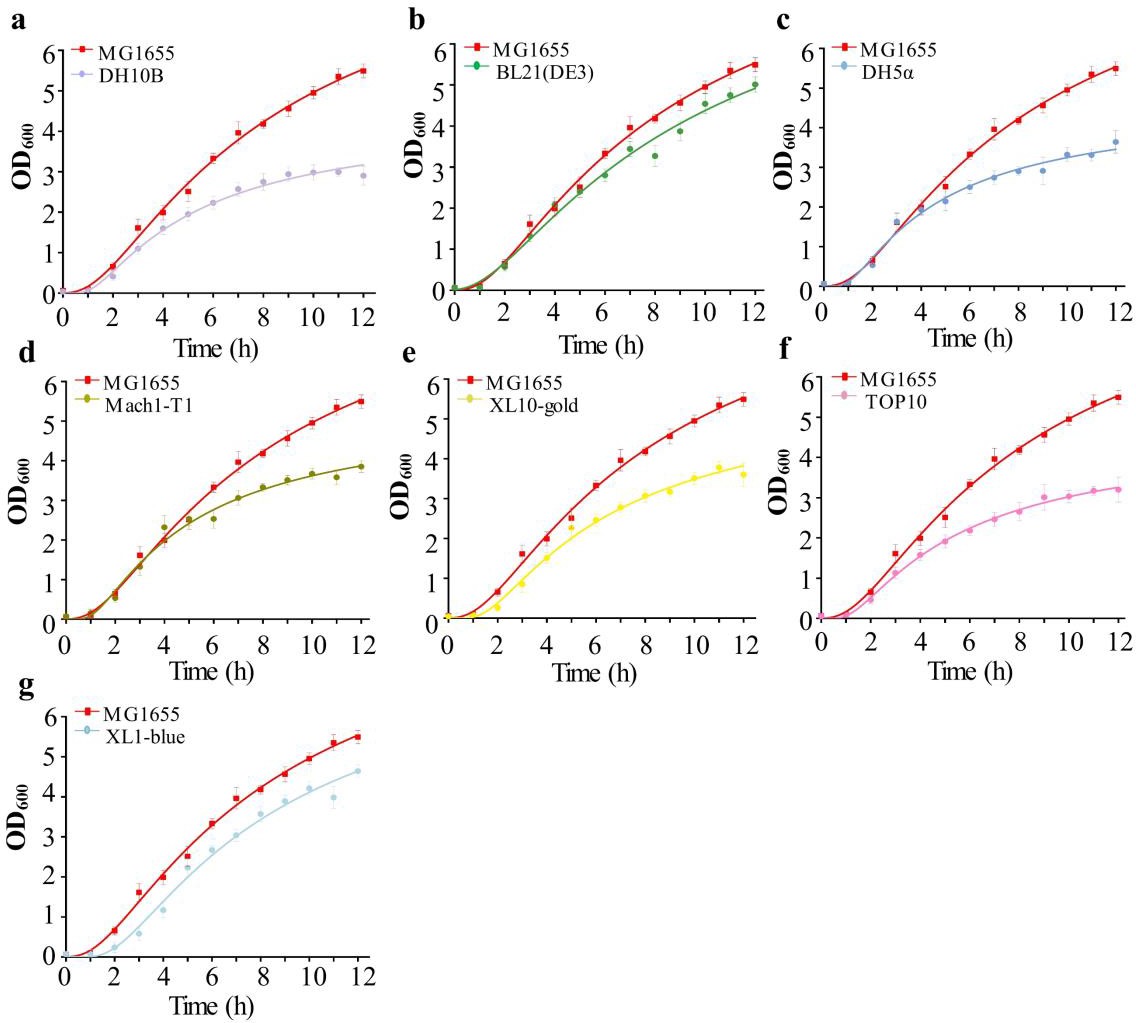

**Figure S1. The growth curves of all eight strains cultured for 12 h.** The curves were plotted by the model of Growth fitting.

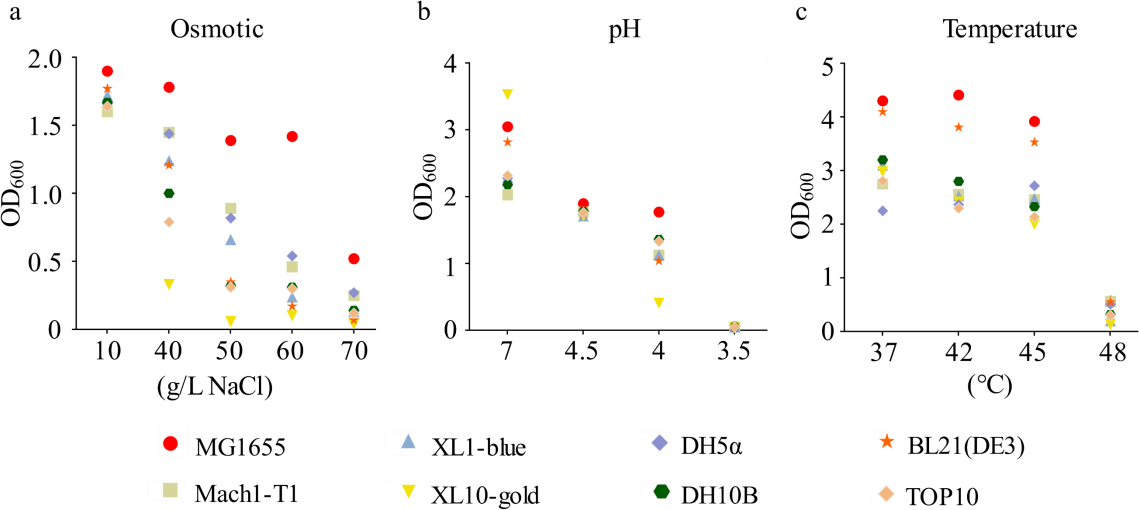

**Fig S2. The maximum tolerability of eight strains under osmotic-, acid-, and heat- stressed conditions.** (a) The OD600 values of eight strains cultured under different

osmotic-stressed conditions for 12 h. (b) The OD600 values of eight strains cultured under different acid-stressed conditions for 12 h. (c) The OD600 values of eight strains cultured under different heat-stressed conditions for 12 h.

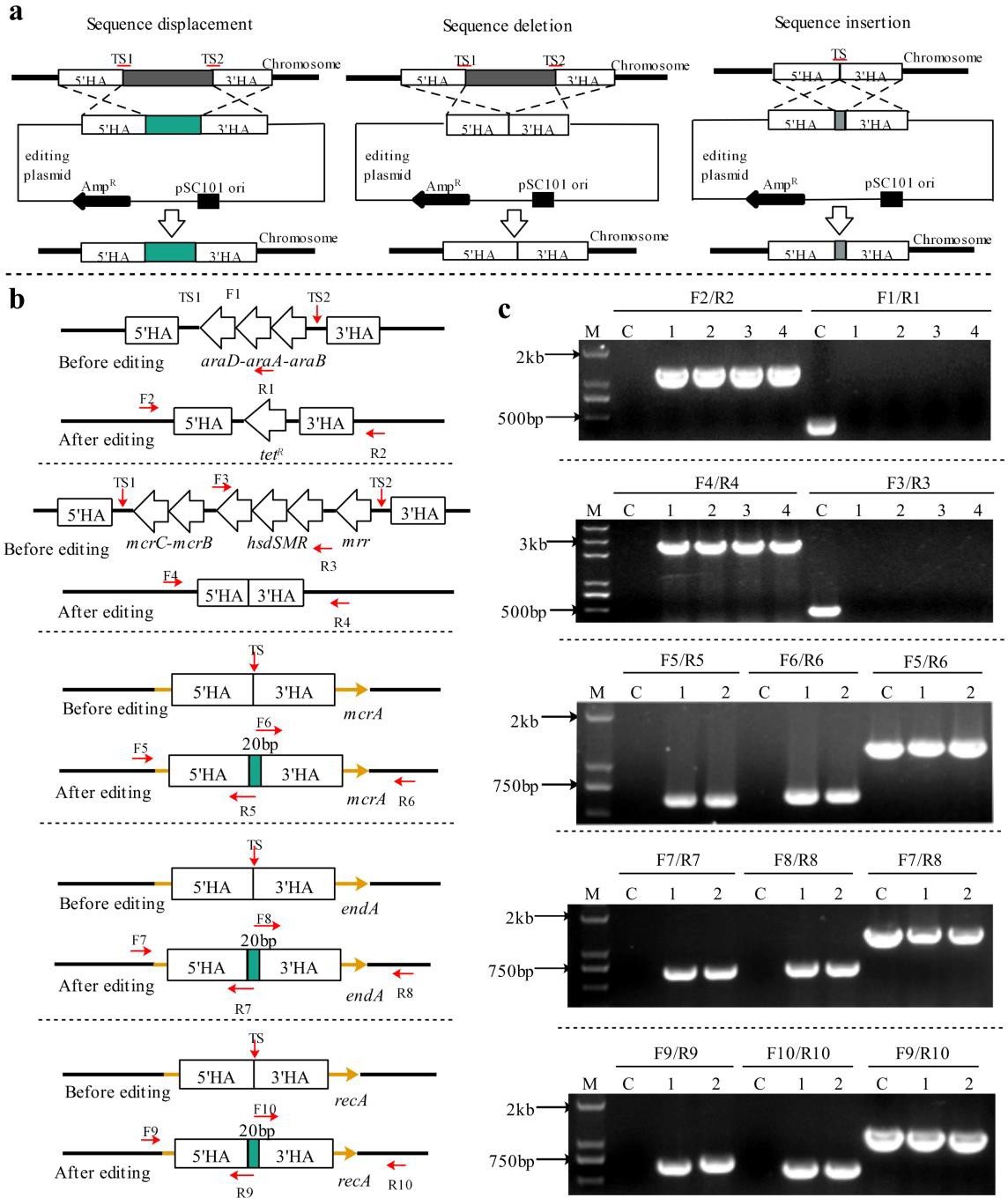

**Figure S3. The principle and results of five rounds of iterative genome editing.** TS, target site. HA, homologous arm, used as donor DNA to recombine with chromosome. 5’HA, the DNA sequence upstream of the target sequence. 3’HA, the DNA sequence downstream of the target sequence. Fn/Rn, a pair of primers used to identify the genome

editing. (A) Three different types of genome editing. (B) Five specific gene editing, insert a 20 base pair sequence to inactive the genes *recA*, *endA,* and *mcrA.* (C) The corresponding gel picture to the B.

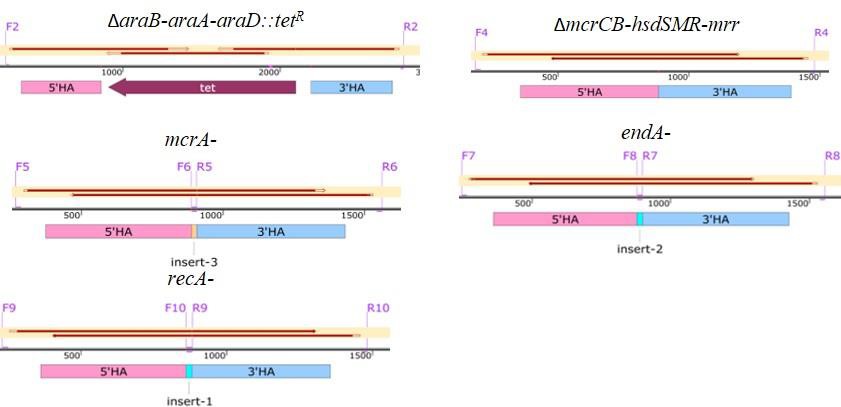

### Figure S4. The sequencing results of the five iterative genomes editing.

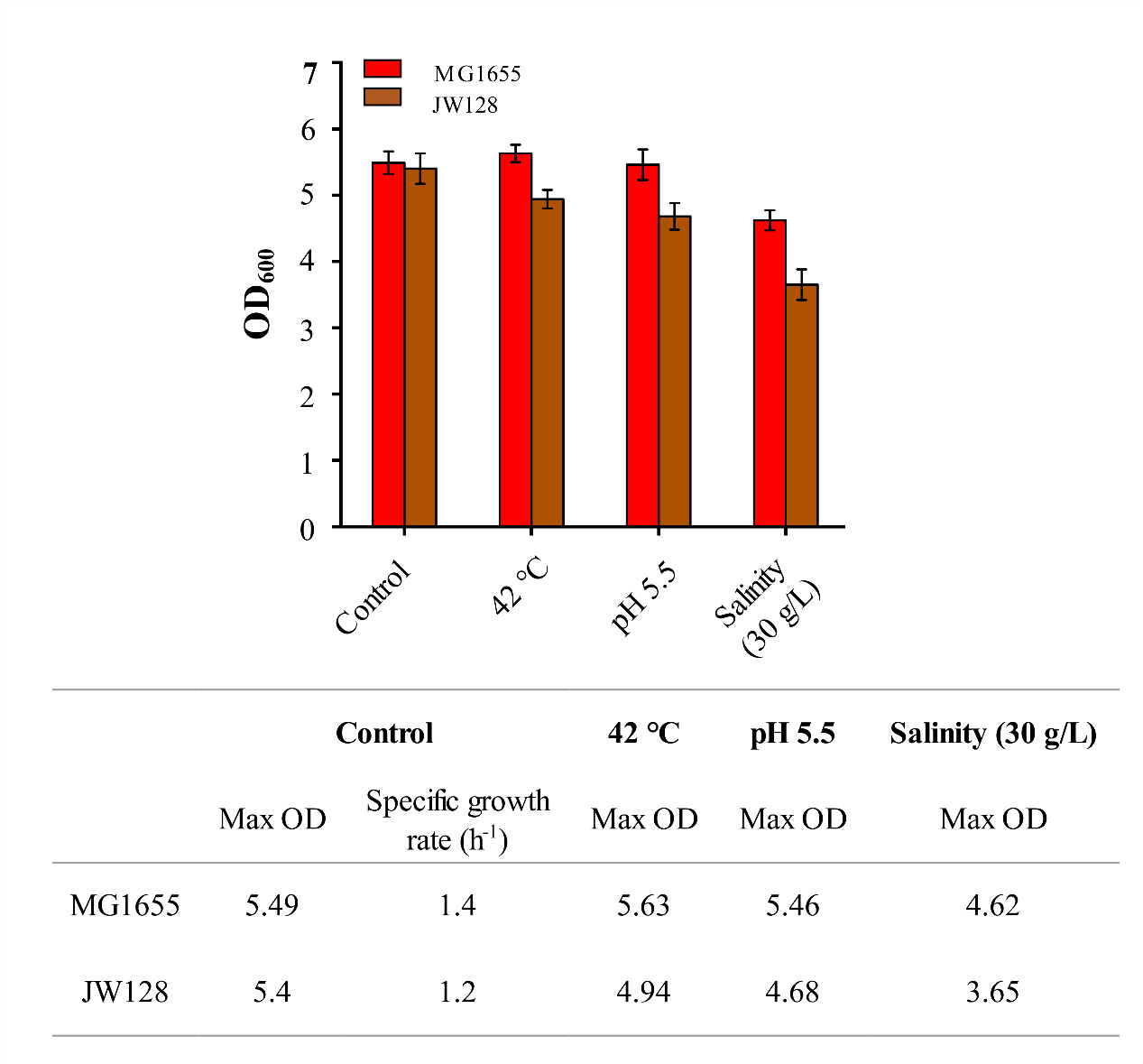

**Figure S5. The growth comparison between MG1655 and JW128.** The OD600 of MG1655 and JW128 cultured under different environments for 12 h, the control environment is LB, 37 ℃, pH 7. Error bars indicate s.d. (n = 3).

### Unit 1. The detail genome sequence before and after five gene editing

The detail genome sequence before and after five gene editing were showed in next section, the sequence highlighted in yellow is the deletion part in the gene editing, the sequence highlighted in green is the sequence of *tet* gene, which was replaced with the gene cluster *araB-araA-araD.* The sequence highlighted in red is the 20 bp‐fragment,

which was inserted into the gene to inactive it.

1 Δ*araB-araA-araD::tet^R^*
  1.1 Before editing
  1.2 after editing
2 Δ*mcrCB-hsdSMR-mrr*
  2.1 Before editing
  2.2 After editing 3 *mcrA*-
  3.1 Before editing
  3.2 After editing 4 *endA*-
  4.1 Before editing
  4.2 After editing 5 *recA*-
  5.1 Before editing
  5.2 After editing
1 Δ*araB-araA-araD::tet^R^*
  1.1 Before editing ccaggcgagtaccgatataggtcagcaagccattttcttcttttacttcttcgacttgcatctgccagccgtcatggctggtaat ggtatcaccagggttgaacatcacgcgggtcacgggggaatcactgcgtgcgtacagacggttttcaccagtagatggga aaagtaaagtgacagttcgcgcatccaccgcgacaacggttccaagtcccaattcgctttctgtatcgctgatccagcgttga ccaagtgtaaaaggcatatgtgttcggctctatatctttaattgcaggcaataaccacccgctaccgtgcttatgaggtagtgg tgttattcaggtccaggaatggaaagggcgctatggtactggatggcaaagcattcgtcacgcatcaaaatggtatctggcg aactcttttttttgctcaaaatagcccaagttgcccggtcataagtgtagcaaaattatcctcaataaaagggagtattccctcc gccacgggttgtagctggcgggtcagatagtgttcgtaatccagtggtgaacgttggtagtccagcggctccgggccgttg gtggtccatacgtacttaatggtgccgcgattctgatattgcaaggggcgaccacgcttttggttttcttcatcggcaaggcga gcggcgcgtacatgaggcggcacattacgctgatactcgctcagcggacggcgaaggcgtttacggtaaaccagtcgcg catccagttcacccgccatcagtttgtcgatggtttcgcgtacatattcctgatatggctcgttgcggaagatgcgcaggtata gctcctgctgaaactgctgggccagcggcgtccagtcggtgcgcacggtttccagccctttaaacaccatccgctgcttgtc gccctcctgaatcagtccggcataacgctttttactgccggtatcggctccgcgaatggttggcatcagaaaacggcagaaa tgggtttcatactccagttctaatgcgctggtcagccgttgtttttgcagcgtttccgcccaccaggcgttaacgtgctgcacca gtgcacgaccgattttcgccgcttcttcttccgaatgtgcgcctttcagccagacaaacgttgagtcggtatcgccgtagataa cgtcgtagccctgtgcttcaatcaacgctttggtttgccgcatgatctgatgaccacgcatggtgatcgacgatgccagccgc ggatcgaagaagcggcaggcggtggtgccgagcacgccataaaaggcattcatgatgattttcagcgcctgcgacagcg gtttgttaccctggcgtttggcttcatcgcgcccgtgccagatgttagtcacaatctccggcaggcaatgtttttctcgcgaga accaggcatcgagaaaaccttcggtactgtgctctggatcaggctgcgccatgccttccaccagcccgacgggatcaatca gaaaggtgcggatgatcgacgggtacaggcttttatagtccagcaccagcactgaatcataaagccctggccgtgaatcca tcacgtagccgccagggctggcgtgcggcggcacttcgccgagattaggcgcgacataaccagcgcgatgcattcgcg gaaaatagagatgaccaaatgccgccaccgaaccgccgtgtcggtccaccggcaggccgttcaccgttgcccgttcgagt aaaaatggcatgatttcagttttgtggaagatctgcgtcaccagctcgcaatctttcaggttataagttgccagcgcaggtttat cttcggcgaaacggcggtcaatttcgtccattcgatcccacgggttatcgatagattttccttcgcctaatagctcctgagcga cagtttccagcgagaatgaagagaaattccagaacgcggatttcagcgcctcgataccgtcgataattagccgacctttagc ctgggcaaaaaagacgccgtttttaaagccgtgctcgcgccactccagctcgctattatcgcgcccaagacgcagcggaa gacggtaacgctcggcatgtttttgcagcattcgcagatcgaactgcaccacgttccaaccgatgatcacatcaggatcgta gttggcaaaccaggcgttgagtttttccagcaactgcgggcggctggcgacgtattccagttcgaaatcaagcgaggagg

cgtcgccattctccggccccagcatataaacgatgcgctgcccgcagccttccaggccgatgcagtacagctcaccgtgg cgggtggtttcaatatctatagaaacccacttgagcggcggacgatagtcgggatgcggtttcagacgggcattaacgata gtgccattgtgcatatcaccctcgacccacaccggtgaggtgataaaccgctccatcagatagcgttctggcggacgcaca tcggcctcgtagacggtaacgccaccttcacgcaggcgcttttcgtaattcatcaattggcgatgggcgcgacagtaaagg ccatacaccggctggcggtgaaaatcctttaacgccagcggtgtcaggcgaaagccttgttcaccctgcaaaatatgctga gcgcggggaacctgatcggcgggaataaacgccacggactcttgcggtgcaagcgtaacctgcaacggcccgttgtccg tcgccagccagaaggagacttctgtcccttgcggggtgtcccgccagtgtcgggttaagataaaacctgcctgcgccacg ctgaaaatccatcaaaaaaccaggcttgagtatagcctggtttcgtttgattggctgtggttttatacagtcattactgcccgtaa tatgccttcgcgccatgcttacgcagatagtgtttatccagcagcgtttgctgcatatccggtaactgcggcgctaactgacg gcagaatatccccatataagcgacctcttccagcacgatggcgttatgcaccgcatcttcggcatttttgccccatgcaaacg ggccgtgggaatggaccagaacgccgggcatttgcgctgcatcgataccctgtttttcaaaggtttctacgatgacgttacc ggtttcccactcatattcgccgttgatttctgcgtcggtcattttgcgggtgcagggaatggtgccgtagaaatagtcggcgtg ggtggtgccggttgctggaatcgactgacccgcctgcgcccagatggtggcgtggcgcgagtgcgtatgcacaatgccg ccaatggaggggaatgcctgatagagcagccggtgagttggcgtgtcggaggagggctttttcgtaccttcaaccacttca ccggtttcgatgctaaccacgaccatatcgtcagcggtcatgacgctgtaatcgacgccggaaggtttgatcacaaagacg ccgcgctcgcgatcaacggcgctgacgttgccccatgtgagcgtgaccaggttgtgttttggcagcgccaggttggcttcta atacctggcgtttgagatcttctaacatgttgactccttcgtgccggatgcgctttgcttatccggcctacaaaatcgcagcgtg taggcctgataagacgcgccagcgtcgcatcaggcgttgaatgccggatgcgctttgcttatccggcctacaaaatcgcag cgcgtaggcctgataagacgcgccagcgtcgcatcaggcgttgaatgccggatgcgctttgcttatccggcctacaaaatc gcagcgtgtaggccagataagacgcgtcagcgtcgcatcaggcgttacataccggatgcggctacttagcgacgaaacc cgtaatacacttcgttccagcgcagcgcgtctttaaacgctggcaggcgtgtgtcgttatcaatcaccgtgatttcaatgtcgt gcatctcggcgaattggcgcatatcgttgaggttcagtgcatggctgaagacggtatggtgcgcgccaccagcgaggatc cacgcttcggaagcagttggcagatccggttgcgctttccacagcgcattcgccaccggcagtttcggcagggagtgcgg tgttttcaccgtgtcgatgcagttaaccagtagacggtaacgatcgccgagatcaatcaagctggcgacaatcgctgggcc ggtttgggtattgaagatcaggcgggcaggatcgtccttaccaccaataccgagatgctgaacgtcgaggatcggtttctctt ctgcggcgatcgacgggcagacttccagcatatgggagccgagcaccaggtcattacctttctcgaagtgataggtgtagt cctccataaaggaggtgccgccctgcagaccggttgacatcaccttcatgatgcgaagcagggcggcagttttccagtcgc cttcgcccgcaaagccgtaaccctgctgcatcagacgctgtacggccagaccaggaagctgtttcagaccgtgcaaatctt caaaggtggtggtgaacgcgtggaagccaccttgttccaggaaacgcttcatccccagctcaatacgcgccgcttccagc

acgttctgtcgttttttgccgtggatttgtgtggcaggcgtcatggtgtagcagctttcgtactcatcgaccagcgcgttaacat cgccgtcgctgatggagttcaccacctgcaccagatcgccaaccgcccaggtattgacggagaaaccgaacttgatctgt gcggcaactttatcgccatcggtgaccgccacttcacgcatgttatcgccaaatcggcagactttcagatgacgggtatcct gtttagagaccgcctgacgcatccaggagccgatacgctcatgggcttgtttatcctgccagtgaccggtaaccacggcat gttgctgacgcatacgcgcgccaatgaagccgaactcgcgaccgccatgtgcagtctggttcaggttcataaagtccatatc gatactgtcccacggcagcgccgcgttgaactgggtgtggaattgcagcaacggtttgttgagcatggtcaggccgttgat ccacattttggccggggagaaggtgtgcagccacaccaccagaccagcgcaacgatcgtcgtaattcgcgtcgcggcaa atagcggtgatttcatccggcgtggtgcccagcggtttcaacaccagtttgcagggcagtttcgcttccgtattcagcgcatta acgacgtgctcggcatgttgggtgacctgacgcagggtttccgggccatacagatgctggctgccaatgacaaaccacac ttcataattatcaaaaatcgtcattatcgtgtccttatagagtcgcaacggcctgggcagcctgtgccggggcggaagttgga agatagtgttgttcggcgctcatcgcccattgctgatagcggcgataaagctgttcaaagcgttgtgcctgctcgctgcacgg ttgcagggttttctctaccgcactggccattttttgctgagctgatgggatgtctgcgtgcactttcgcggcgacggcagcaaa aatcgccgcaccgagcgcacagcactggtcagaggcaacaatttgcagcgggcgattcagcacgtcgcagcaggcctg cataatgacctggtttttccgcgcgatgccgcccagtgccatcacgttattaacggcgatcccctgatcggtaaagcactcca tgattgcgcgtgcgccaaaggcggtggcagcaatcaaaccgccgaacagcagcggagcgtcggtagcgaggttaagat cggtaatcacccctttcaggcgttggttagcgttcggtgtgcggcggccgttaaaccagtcgagcaccaccggcaggtgat ccagagacggatttttggcccatgcttcggtcagcgccggaagcagttgtttctggctggcgttgatttgcgttttcagttccg gatgctgggcggcaagctgttccagcggccagccgagtacgcgaccaaaccaggcgtagatatcaccaaacgccgattg gcctgcttccagaccgataaatccaggcaccacgctgccatcaacctgaccgcaaatacctttaactgcccgctcgccaac gctctgtttgtcggcaatcagaatgtcgcaggtggaagtaccgataacttttaccagtgcgttaggctgtgcgcctgcgccaa ctgcgcccatatggcagtcaaacgcgccgccggaaatcaccacgctttcaggcaggccgagacgctgcgcccattccgg gcataaggtgcccaccggaatatcggcagtccaagtgtcagtgaacagcggggaaggcaaatggcgattgaggatcgg gtccagctcatcaaagaaactggctggcggcaggccgccccagctttcgtgccacagagatttatgcccggcgctgcaac gtccgcgacgaatatcctgcgggcgggtggtaccggaaagcagagctggcacccagtcgcacagctcaatccacgatg cggcagattgcgccacggcgctgtcctggcgagtcacatgcaggatttttgcccagaaccattcgctggaataaataccac caatgtagcgggagtagtcaacgttgcccggcgcgtggcacaaacgggtaatctcttccgcttcttcaaccgcagtgtggtc tttccacaatacgaacatcgcgttcgggttttcggcaaactccgggcgcagcgccagcacgtttccgtcggcatcaatcggt gcgggcgtcgagccggtactgtcaacgccaatcccgaccacagctgcgcgctgttcgacgctaagctctgcaagcacgg ttttcagtgccgcttccattgactcaatgtagtcacgcggatgatgacggaactggttattcggggcatcacaaaattgcccttt

ctgccaacggggataccactctacgctggtggcgatctcttcaccggtagcgcagtccaccgccaaagctcgcacagaat cactgccaaaatcgaggccaattgcaatcgccatcgtttcactccatccaaaaaaacgggtatggagaaacagtagagagt tgcgataaaaagcgtcaggtaggatccgctaatcttatggataaaaatgctatggcatagcaaagtgtgacgccgtgcaaat aatcaatgtggacttttctgccgtgattatagacacttttgttacgcgtttttgtcatggctttggtcccgctttgttacagaatgctt ttaataagcggggttaccggttgggttagcgagaagagccagtaaaagacgcagtgacggcaatgtctgatgcaatatgg acaattggtttcttctctgaatggtgggagtatgaaaagtatggctgaagcgcaaaatgatcccctgctgccgggatactcgtt taacgcccatctggtggcgggtttaacgccgattgaggccaacggttatctcgatttttttatcgaccgaccgctgggaatga aaggttatattctcaatctcaccattcgcg

1.2 After editing ccaggcgagtaccgatataggtcagcaagccattttcttcttttacttcttcgacttgcatctgccagccgtcatggctggtaat ggtatcaccagggttgaacatcacgcgggtcacgggggaatcactgcgtgcgtacagacggttttcaccagtagatggga aaagtaaagtgacagttcgcgcatccaccgcgacaacggttccaagtcccaattcgctttctgtatcgctgatccagcgttga ccaagtgtaaaaggcatatgtgttcggctctatatctttaattgcaggcaataaccacccgctaccgtgcttatgaggtagtgg tgttattcaggtccaggaatggaaagggcgctatggtactggatggcaaagcattcgtcacgcatcaaaatggtatctggcg aactcttttttttgctcaaaatagcccaagttgcccggtcataagtgtagcaaaattatcctcaataaaagggagtattccctcc gccacgggcatcttggagtggtgaatccgttagcgaggtgccgccggcttccattcaggtcgaggtggcccggctccatg caccgcgacgcaacgcggggaggcagacaaggtatagggcggcgcctacaatccatgccaacccgttccatgtgctcg ccgaggcggcataaatcgccgtgacgatcagcggtccaatgatcgaagttaggctggtaagagccgcgagcgatccttg aagctgtccctgatggtcgtcatctacctgcctggacagcatggcctgcaacgcgggcatcccgatgccgccggaagcga gaagaatcataatggggaaggccatccagcctcgcgtcgcgaacgccagcaagacgtagcccagcgcgtcggccgcc atgccggcgataatggcctgcttctcgccgaaacgtttggtggcgggaccagtgacgaaggcttgagcgagggcgtgca agattccgaataccgcaagcgacaggccgatcatcgtcgcgctccagcgaaagcggtcctcgccgaaaatgacccagag cgctgccggcacctgtcctacgagttgcatgataaagaagacagtcataagtgcggcgacgatagtcatgccccgcgccc accggaaggagctgactgggttgaaggctctcaagggcatcggtcgacgctctcccttatgcgactcctgcattaggaagc agcccagtagtaggttgaggccgttgagcaccgccgccgcaaggaatggtgcatgcaaggagatggcgcccaacagtc ccccggccacggggcctgccaccatacccacgccgaaacaagcgctcatgagcccgaagtggcgagcccgatcttccc catcggtgatgtcggcgatataggcgccagcaaccgcacctgtggcgccggtgatgccggccacgatgcgtccggcgta gaggatccacaggacgggtgtggtcgccatgatcgcgtagtcgatagtggctccaagtagcgaagcgagcaggactgg gcggcggccaaagcggtcggacagtgctccgagaacgggtgcgcatagaaattgcatcGaacgcatatagcgctagca gcacgccatagtgactggcgatgctgtcggaatggacgatatcccgcaagaggcccggcagtaccggcataaccaagcc tatgcctacagcatccagggtgatggtgccgaggatgacgatgagcgcattgttagatttcatacacggtgcctgactgcgtt agcaatttaactgtgataaactaccgcattaaagcttatcgatgataagctgtcaaacatgagatcagatccttccgtcgaggc caattgcaatcgccatcgtttcactccatccaaaaaaacgggtatggagaaacagtagagagttgcgataaaaagcgtcag gtaggatccgctaatcttatggataaaaatgctatggcatagcaaagtgtgacgccgtgcaaataatcaatgtggacttttctg ccgtgattatagacacttttgttacgcgtttttgtcatggctttggtcccgctttgttacagaatgcttttaataagcggggttaccg gttgggttagcgagaagagccagtaaaagacgcagtgacggcaatgtctgatgcaatatggacaattggtttcttctctgaat ggtgggagtatgaaaagtatggctgaagcgcaaaatgatcccctgctgccgggatactcgtttaacgcccatctggtggcg

ggtttaacgccgattgaggccaacggttatctcgatttttttatcgaccgaccgctgggaatgaaaggttatattctcaatctca ccattcgcg

2 Δ*mcrCB-hsdSMR-mrr*
  2.1 Before editing gatttcgctgatcagtggtttgctgctgcgctgggcaggccggtagaactgacggtgttcgactcgccgcgcgatatccctc accatcgtaaactgacggtgacttttgaggatggtcaggtattgaagatccgcttcgatcaggggatgggctactggcgcat caacttttcatcgcaatggcattactttgatttccgcgatgacgtttctttccagttagtcaaaatggctcaggcctgcaaggaa gggaatgtcgccaacagcgaagagagttgggcaacggatgtgctggtggaggtgatcgcctcctgatgatgagccgctc ccgatgtggtgtcgggagcggtattttctataaaacttaccgcttatttgagatattcatcgaaaatgtcgagtaattcttgatgta tacacggccattcctgacctaaattgacggtacacaagccaatatcgaagccattaattttataacgatgtttcactgcggtatc tacgtggggatatattaataacccccctatgttttcgccattttcaggctttaacgaccataagtaattcatcagttgataaagatt ttgcgaatgaaatttttctgttcccattcgtcgtgaaaaaatgctcttatagtatttggcgtcaacgataagtattttttctgatgag

cgaatggtgatgtcagtttccattcgaggtaacaaattaagtgactgatccgatatactcgatgcatcccattttaaataagagc

gggttgtgtttgcagacgttaattcacgacggcaaaattcataaagaaacttttgataaagtaatgacatctctttttcgtttctttc

aaaatcatagaaacggtagtgtcctttgttttgacctggaatagaattattgacgatgaatttgcagacactgataacgaatttat

aataacgcgtattttttccgccattcagatagctgaaatgctgcggagttaaatgaagagtgctaatgcccggtaattttctata aagtgaacgagcttcatctctgatagttgaatttaacttttcatgcttaattaatatggctaatgtgctttttataattcggttagcca gcgtgtcttcattaagcatatcaaaagtactgacggttttcccatgattaagatggaagccgcgtattgttttagcaaactctatt cgccctttgatgccaggaatgatctcggtgttaggattgtaatcaagctcaagccctcggcgtgaaagctgtaaaaccccttt

atttaatacataccccaggatatcaagaagattgttaccgggtatggcttcaaggtttgcctgcttaatttcctgtaaataacccc

atgcataggtaagcatgtaatagatattacggacaggtatcacgggctgttccactatgagtcccctaataatttgttggtccat

ttctgttgtttataggggtcatcaaagaaatattcttcgagtaaaggggcgatatccgtcatcacaatttcattaagccattgcgt atccggagaggtgccatcttccaacccacagcagaagtaactatgcccaatgcggaatcctttcccaaggatagtggcctct

ttgctgatttcctggttcaactcgttcattttttggcataaagactcaacaaatgaaggttctgcttttttattcagtaaaaaattccg

gaactgtggtgtatcaaaacctggctcaatatctatgaaagaaaatcgtctgcgtagggcatagtcaacaacggccagaga gcgatcggcagtattcattaaaccgatgatataaacattctccgggacatagaatcgttcttcatcgttttcggagtaggttagg

ggaacagaccagttttcacctcgtttatcatgttccattaacatcatcacttcgccaaatactttactgagattggcacgattgatt

tcatctataataaaaatatactttttctctggctgctctttagcttgctgacaaaaattgtaaaatatgccgtctttacgtcggaagc

cgacgccattcggacgatagccctgtataaaatcctcatagctataagattgatggaactgaaccatattgacgcgttgcgga gccttttctcctgtcagcaagtaagccagacggcgtgcaacaaaggtttttccaacgccgggcggcccctggaggataata

ttttttttgatggttaatcgtttgagtatcgtctctattgtggtttcagggataaacaaatcatttaacgcatcttccagacagtatga

ttcagtttttgacataggtggaataacactcttgccagaattaaatattaatttatagtcgttgattatgttgtccagcatagaggc aaatcgggtgtaatcaataccctgtgagactttttgggaacaggcgtaataggactgtccgtattttttaggatatacacccga agttgcctgaaaatactctgcgattgttttaggtatgtctgaagagaactgccattgggcatgtggttcattcgtgtcgcttatac cataagccaaaaccaactcatcaaaatctttataatagagaataacgggatatataccgttagaagcttcctgaccttctccaa gaaatgcaaaccagggaatagacgtaaaattaccataaccgaaactcaattttactcgcaggttacggtaagacgttggata atctttagtggattgcgaacgttgttgctgtgcttgcttaataaatttttcaatccagggttgaatagattccataagatatgccttc ctcattgctaagcctctattatcgctttcgcaacgtactgaaacaatagatttttactgcaaaatcagactggtaaatatttactga gggggaaagtttctattgagtcagtggaaggctcccggtggttaaccgggagtaaacgctgttacgcgactttctgtttaccg gcaatcactccaataaacgcctgcacctgcttttgtttacgcgccgacagtttgcacacctggcgtagcgactgcatcagttc gctctcctcggcggcgggtggttgggcggtgaggacaatacagccttccatcactttgacatctaccgccgtgccagtggc aaaaccggcggcttccagccactgacctttcagggtgatggcgggaatacggctgtaatccgggtagcgactcgcataac cgacggtgacatgacggttatttgccggggagacttctgcttcgaacggttgtgcaatagaatgcgtgtcagtcataactgct attctccaggaatagtgattgtgattagcgatgcgggtgtgttggcgcacatccgcaccgcgctaaatacctgtatatatcatc agtaaatatggggaaagtccagctaaaaatagaataaaatgggcaatttctggaatgatttaaatatatttatgtgggttatgatt ggcgtgaaataataaaaagcgcaccggaaaggtgcgccagaaaataatgttcaggattttttacgtgaggcttttttaccccc gctagctgcgcgttcagctttgattttttccagcaacgcggcggcgctgttttctccgctgatcaaatccgggttttcggcccg ccactgggcggtaagttcaccacggaacgcttttgccaggatggattgcgtcaggttgttgacgcgggctaaggcgttgttg acctgtttttctatggtgtcggcgtaggcgaagagttgctcgacgcggcgaacgatttcggcttgttcttttactggaggtaata aaacaacttgggatttgatatcttttcctgaaatacctttttgaccagaagttgttttcacgcagttcatcattgcatttcgtgctga gggggatgaaaaaaatatttcgatatattctggtaaagcatctttggttaatcgagctcgaataagtttatcaggatatagcaaa ttttgatgttgtaattttttcaataacccacaaacaccaacaaattctaaacttccgttatagcgagtaaataaaagatctccatctt gtaatttgtggcggtttagttcactttctgaacattctagaaaccgaatatcgttttgatctacatggccagcacgtacagaacta atgcgtagtattggatgaccaacaccactttcatttggctttgatgaaagaccattacgtaattcagttaagatagattcaaaatt taacttcttaaatacagaatgttgcggctcaaaattacgccatttttctgtcaattttccattaactgcgccccccaataccgcttg acgaaaacgtttcaggatttgtgggatttgctcaaaacgtgctttggtgctgtctacctgcgccagcagcgtatcgagtttttca gcgatgattttttgttcggcaagtggtgggattggtatatttatcaaatcaaagcttgccggcttaatattattaatatttgcacca gcagaaagtgatgaaattttgtttcgataaagagaagattttgtgaaatgagcaataaaaccagaaaatataagtttttcagga cgtaatacaccgcaaaatgcgccgaaactacattcaaatggtagatgctgatgtgcggatttaccaactacggatttgctccc tgatgacattgcaataacaatatcttcaggagatattttttgactttctttaacaagatttttaggaacaaaaaccaagtccgtagt

atcaaacttgccattctgaatattgttcgcacggataagaggcaaataatcatcttttagataatttattgcctgctcttttttatacg ttactcctcggattagagttgtgaccgtagatactggggcgataacccacccctccggcaatttccccgcactcattccttcac cccaccaaacgcttcttccagcaactgacgctgcaaatcggcctcatcgctcgcccccagttcacgcatcagcgcatccag ttcagacagcgcctgtaccagttcgcccatcgcttctgccgctaatacatccggctccggcaggctgtcggcatcaatactgt ctttatctttcagccaggagatatccagcgaatcggattttgcggtgcggatccactcacggctgaacttgcgccagcggct ggtagcaagatgctggtcggtgtttttgttctcttcgctgtcggcaacttccgtctcttcggcgttaaaactccattcaccttcagt gcgcgggcttaaaccgtgcgggtcttcgccatacacgcgctcaaacggctgcaaatgctcgtcggtaaacggtgtgcgctt gccgaaactcggcatattggtacgcaggtcatacacccacacatcatcggtacagttcttatcctgattcgggttcgccaccg tccctttggtaaagaacagcacgttggtcttcacgccctgagcgtaaaaaataccggtcggcagacgcagaatggtgtgca gatgacacttatccatcaggtcacgacgaatgtcggtgcctttgccgccttcaaacagcacgttatccggcaccaccaccgc cgcacgaccgccgggatgcagcgtttcgataatatgctgcataaagcacaactgtttgttgctggtcgggtgaacaaaggt gcgggtaatgttggtgcctgcggcgctgccaaacggcgggttagtggcgacaatatgcgccttcggcaggttttcaccgtc gctacccagagtgttgcccagacggattgcgccgccgtggtcgaggttgccttcaatatcgtgcagcaggcagttcatcag tgccagacgacgggtgccgggcaccagttcgaggccgataaacgcgcggtggatctggaaatcctgcgtgtcgccatca aggtcgtccagatcattggtttgcgacttaacatagcggtcggcttcaatcaaaaagcccgccgtacctgccgccgggtcct gcaccacttcacgcggctgcggtttcagcagatgaataatggttttaatcagcggacgcggggtgaagtactggcctgcac cagacttggtttcattcgcgttcttctgcaacagcccttcgtacatatcgccgaagtcatcgcgcgacttaccgtgcgcgccgt tgtaccagtccagcgaatccatattgctgaccagtgcggttatttgtttcggctcggtgatggtggtactaacattatgaaaaac tgcctgtaccagctttttgtcatcttcgcctaaatgcacgagcatttttcggtagaactgcaactgctcctggccgatgcgggat ttcaggtcatcccagcggtaaccttccggcaggtattccgcttcctgaccggtctctttacacattttcaaaaacagcagcgag gcgagttcattgacgtagttttgataggaaacgccgccatcgcgcaggttgtcgcacagcttccacagcttcgcgaccagat cgttattgttcattgtgagttccgtaaattaagcagcggcccaaattcatcgagccgcagaagaagaaattgccgagggtaat atacacaaaatcattcaggttgcatcaaggcggcaagtgagtgaatccccgggagcgtacagaagtacgtgaccggggtg aacgagcgcagccaacgcagaggcagcctgaaggatgaagtgtatacgtgtcaggccagctcgtcccagatataatcgc tgaatttgcccagcagggtatcgagattatcgtcaaaggttctttgcagcatcgccttcccgccgcgacggtggaagttgcc ggttttgaagacatcgtcgtcgagcaccactttctctttcagcgcctgcgctaaacgatcgagccagcttaattgctcgctgct ccagtcgttttcgcccttaatgcgcgtcagcgcgtgatcgacacgttcctcaaacggtttcagcgcatcgcccaccgcagcg cggcgaatatgaccaatcagccgggcggcgatatcttcattgcgcgtctctttccatgctttgcgcagggaagattcctcaaa gtgctggcggtcaaaccactcctgtagctcgaccagccctttacgggtgagatcgcgcgggcgattaataactgcctgcaa

tgccggttgcgcgttcggggaacgttgcaccagcgagtcaaaggcttcgaggaaatcctgcggcgtgtcgtaatcaccgt acagcgattttacactcaccacttcatcgtcgatatcgaggaagatcggcgcatcattcaggttgttgatgtccgttttcagcttt tccagacgggcgataaagccaggcagtttgttaaagacttcggcgctccagtgcggccctttttcccgcaggcgcgaggc gaagccgttaaagttcacgcccgccgcgtcctggcatagctcatccagacgacgcacctgtttatctatcgtttcgctgcggt cacggttaaacgtggccagaccgatgatacgctggagcttcgccaccagttgttcatggctgtgctcggcaaaactgcggc catccgcttcggtgattttataggtttctgaatcggtaatttcattgaccagcgtttgcagttccaccttcgggcgcaccaccac cggacgcatggtgtcgacgctctccagcgtgctgtagatatcgacacagtcaaaaatcttaaagctggttttattcacctccg ggcataagcgcgtggcgcggcctttcatctgttcgtacagaatgcggctgcgtactttacgcaggaacacgatattacagat cgacggaatatcgacgccggtcgtcagcaggtcgacggttaccacgatattgggcagccgctctttattgaagcgggtgat catggtctgcactttgcgcgcgtctttatcggcatcaccggtgatcttgatgatcgcgtcgtgctccagttgcggatactttttct tgaacgcggcacgcagctcttccaccaccatatcggcatgggcattggtgacgcagaagaccagcgttttttgcgatccgg tcgggtcaagataattggtgagttcgttacagacggcgcggttaaacgccgggatcaccaggccacggttaaagtcggcg acttcaaaatcctgatcgtcttccagggtgtcattgatcacttctccctgcgggctgatgcgctctacctgctcgcctttggaga gataaaccccctcctgcgcgttgcgggtgatgatctgaataggcggatcctggtcgatcagaaaaccgtcgataaccgcg gtacggtaggtataacggtaaaccggctcgccgaaaatctgcacagtatgtagcgccggggtggcggtgagagcgattttt accgcatcgaagtgatcgagaatgcgacggtaggcagagacgtaatccagctggctgcggaactgcagttcgccttcggt ctgctctttatcgagaatatagccgcgatgcgcttcgtcaacgacgatacagtcgtaacgggccaccggcatcggttcatct gattgcagggtgcgtttcaccagcgactgtacggtggcaacgtgaattttggtgctgtcttccgggaatttatccgtcagccct ttaatgtcgaaaatgctgttgaaggtgtcgccgttaatacgcgtatcttcaaacgcgcccagcgcctgttcgccaagagaac ggcggtcgacaaggaagagaatgcgtttaaaacgctgggactggatcaggcggaacatcatggcgattgccgtacgggt tttaccggtaccggtcgccatcgccagcaggatctcttgttgccccttgacgattgccttttcaaccgcgcggacggcatcttc ctgataataacgcaggcccagctcgctcatgccagggttatcggcaaaccactgattctgtttttgcggttcgctgccgagca tttccagcagctcttccgggcggtgccactcgggtaaggctttcgacatattgcgggtatcacgcacgtcgcgataccagat gccgcttttggtcttcattgttgcgcggtattcgcgcccgttggtcgagtagcagaaggggattttaaaccgttgtttgccgctg gtgtcctgccagctggtttcatactctggcactgcttcatgcacttcatccggtgagtagtgctcaagcaaggtttcccgcagg aagccattatcgaaacatttactgtagcgatacgactcattgagcctggcgggaacgtcgatattgttacgtttcgcctctacc accgcgatgggtttgaggccgacaaacagcacataatccgcaaagccctgattacccgtttcatcttttccggtcggccattc ggcaatggctttattgacgccgggttccggacgtgcgcctttggagaagcgcagggttttgctgtcggcctgccagcctgct ttacgcagttgcgcatcaatcaggaagcgactctcttcttcgctaaggttgagtgtgcgcttgatggcctgatcggtaatttcttt

gtggtaagccttacgttcctgttcggtctgttttgccagttccgcgttcttctcggcgagctgtgcttccagtgccgcaaggcg agcctgggtttgcgcttcggtttcctgctgtttgccttccagaatggcgatatagccgttcagggcaaccagcttctgctgttgc gcttcgacttctgcctgagtctgcgctttttctcgcacctgctgttcaagctgttgttttagcgtcagcacttcctggtgatagag gttttcaccacgttccggcaacacaaacaccggcaccgggaagtcataatctttagtgaccagacggtagtaccagacagc caggcggaacccgagtcgcaggcacatctgggcatcgttgagatcgttatgatattcgtgcaccgcctggttaccaatgcg gcgtaatttgtgaaatacagagaggatgttgtcatcaacaaaggcgattttgccgagttcacgcaggagatcgtgttgattctc acaaggggggatgttgagtaacagaccaagatgtttcgctgtggcttcgccaaacatacgcattttaatcagcgtcgtgttgg gatcatccgggtagttattttccgccgcacaggcgatggcataagtgaagtcgttgacgcccttcaggaattcaaaattggat ttattcatcattgttattaatccattgctgtgcgggcctgtccaaatatttaaggcccataacatctcatcttagctttctgtaccttt ccgggcaatgaccacggtcacagcaactgactcatttctaacgtgttcgtctatttttgtagtgctatagtagccgaaaaacat ctacctgattctgcaaggatgtactatgacggttcctacctatgacaaatttattgaacctgttctgcgttatctggcaacaaaac cggaaggtgcagccgcgcgtgatgttcatgaggctgccgcggatgcattaggactggatgacagccagcgagcgaaagt cattaccagcggacaacttgtttataaaaatcgtgcaggctgggcgcatgaccgtttaaaacgtgccgggttgtcgcaaagtt tgtcgcgtggcaaatggtgcctgactcctgcgggttttgactgggttgcgtctcatccccagccaatgacggagcaggaga cgaaccatctggccttcgcttttgtgaatgtcaaacttaagtcacggccggatgccgtcgatttagatccgaaagccgactct cccgatcatgaagaacttgcaaagagcagcccggacgatcggttagatcaggcgctaaaagagcttcgtgatgcggtggc tgatgaggttctggaaaacttattgcaggtttctccttcgcgctttgaagtcattgttctggatgttttgcatcgcctggggtatgg cggccaccgtgatgatttgcagcgtgttggcggtactggagatggtggcatcgatggtgtgatatcgcttgataaacttggc ctggagaaagtttatgttcaggcaaaacgttggcagaatactgtaggcaggccagaattacaggcattttacggcgcactgg ctgggcaaaaagcgaaacgtggggtgtttattaccacttctggatttacttctcaggcgcgtgactttgcccaatccgtcgag ggtatggtgttggttgatggggaacgcctggtgcacttaatgatcgaaaacgaagtaggggtttcttcacgtttgttgaaggt gccgaaactggatatggactattttgagtgaaatatcaggccggatgcggctgcgccttatccggcccataaccccttacttc ctcaaccccgcaaacgcagcccgaatctcttcctccggcagctggatcccgataaacaccatcgtgctatgcggtttttcatc gccccacggcctgtcccagtcggcgctgtagaggcgctggacgccctggaacagcaggcggttaggttcgccgtcaatc cacagcatccctttgtaacgtagcagtttatccgccgactccagcagcaggttttccatcacgcgggaaacttcgctgatatct accgggtaatccagttccaccacaatcg

2.2 After editing gatttcgctgatcagtggtttgctgctgcgctgggcaggccggtagaactgacggtgttcgactcgccgcgcgatatccctc accatcgtaaactgacggtgacttttgaggatggtcaggtattgaagatccgcttcgatcaggggatgggctactggcgcat caacttttcatcgcaatggcattactttgatttccgcgatgacgtttctttccagttagtcaaaatggctcaggcctgcaaggaa gggaatgtcgccaacagcgaagagagttgggcaacggatgtgctggtggaggtgatcgcctcctgatgatgagccgctc ccgatgtggtgtcgggagcggtattttctataaaacttaccgcttatttgagatattcatcgaaaatgtcgagtaattcttgatgta tacacggccattcctgacctaaattgacggtacacaagccaatatcgaagccattaattttataacgatgtttcactgcggtatc tacgtggggatatattaataaccccccacgtggggtgtttattaccacttctggatttacttctcaggcgcgtgactttgcccaat ccgtcgagggtatggtgttggttgatggggaacgcctggtgcacttaatgatcgaaaacgaagtaggggtttcttcacgtttg ttgaaggtgccgaaactggatatggactattttgagtgaaatatcaggccggatgcggctgcgccttatccggcccataacc ccttacttcctcaaccccgcaaacgcagcccgaatctcttcctccggcagctggatcccgataaacaccatcgtgctatgcg gtttttcatcgccccacggcctgtcccagtcggcgctgtagaggcgctggacgccctggaacagcaggcggttaggttcg ccgtcaatccacagcatccctttgtaacgtagcagtttatccgccgactccagcagcaggttttccatcacgcgggaaacttc gctgatatctaccgggtaatccagttccaccacaatcg

3 *mcrA-*
  3.1 Before editing Cgtagtatgcggcatcttgtcgtgctggtggaggagttgcgcgaacgaggcatcaactttcgtagtctgacggattcaattg ataccagcacaccaatgggacgctttttctttcatgtgatgggtgccctggctgaaatggagcgtgaactgattgttgaacga acaaaagctggactggaaactgctcgtgcacagggacgaattggtggacgtcgtcccaaacttacaccagaacaatgggc acaagctggacgattaattgcagcaggaactcctcgccagaaggtggcgattatctatgatgttggtgtgtcaactttgtataa gaggtttcctgcaggggataaataaagttaaagacactttgtgtacaaaagaaagtaaaacaacagcaacttgttgcaatttta tcaataaaagtagtattgtcgtgaaaaattgattaaagattaatattatgcatgtttttgataataatggaattgaactgaaagctg agtgttcgataggtgaagaggatggtgtttatggtctaatccttgagtcgtgggggccgggtgacagaaacaaagattacaa tatcgctcttgattatatcattgaacggttggttgattctggtgtatcccaagtcgtagtatatctggcgtcatcatcagtcagaaa acatatgcattctttggatgaaagaaaaatccatcctggtgaatattttactttgattggtaatagcccccgcgatatacgcttga agatgtgtggttatcaggcttattttagtcgtacggggagaaaggaaattccttccggcaatagaacgaaacgaatattgata aatgttccaggtatttatagtgacagtttttgggcgtctataatacgtggagaactatcagagctttcacagcctacagatgatg aatcgcttctgaatatgagggttagtaaattaattaagaaaacgttgagtcaacccgagggctccaggaaaccagttgaggt agaaagactacaaaaagtttatgtccgagacccg
  3.2 After editing cgtagtatgcggcatcttgtcgtgctggtggaggagttgcgcgaacgaggcatcaactttcgtagtctgacggattcaattga

taccagcacaccaatgggacgctttttctttcatgtgatgggtgccctggctgaaatggagcgtgaactgattgttgaacgaa caaaagctggactggaaactgctcgtgcacagggacgaattggtggacgtcgtcccaaacttacaccagaacaatgggca caagctggacgattaattgcagcaggaactcctcgccagaaggtggcgattatctatgatgttggtgtgtcaactttgtataag aggtttcctgcaggggataaataaagttaaagacactttgtgtacaaaagaaagtaaaacaacagcaacttgttgcaattttat caataaaagtagtattgtcgtgaaaaattgattaaagattaatattatgcatgtttttgataataatggaattgaactgaaagctga gtgttcgataggtgAGTGGTTCTATCGAGAGTCGaagaggatggtgtttatggtctaatccttgagtcgtgg gggccgggtgacagaaacaaagattacaatatcgctcttgattatatcattgaacggttggttgattctggtgtatcccaagtc gtagtatatctggcgtcatcatcagtcagaaaacatatgcattctttggatgaaagaaaaatccatcctggtgaatattttacttt gattggtaatagcccccgcgatatacgcttgaagatgtgtggttatcaggcttattttagtcgtacggggagaaaggaaattc cttccggcaatagaacgaaacgaatattgataaatgttccaggtatttatagtgacagtttttgggcgtctataatacgtggaga actatcagagctttcacagcctacagatgatgaatcgcttctgaatatgagggttagtaaattaattaagaaaacgttgagtca acccgagggctccaggaaaccagttgaggtagaaagactacaaaaagtttatgtccgagacccg

4 *endA-*
  4.1 Before editing ccggagccaaaactctcttacacccagcgcggaacctccgccggaacggcctggctggaaagctatgaaattcgcctcaa tcccgttttgctgttggaaaacagtgaagcttttattgaagaagtggtaccgcacgaactggcacatttgctggtatggaaaca tttcggccgcgtagcgccacatggcaaagagtggaagtggatgatggaaaacgtgctgggtgttcccgcccgtcgtacgc atcagttcgaactgcaatccgtgcgtcgcaacaccttcccctaccgctgcaagtgccaggagcatcagcttaccgtacgcc gccataatcgcgtagttcgtggcgaggccgtctatcgctgtgttcactgcggtgaacagctggttgcgaaataaccatctga actatcaggaactttcctgatctggctgattgcataccaaaacagctttcgctacgttgctggctcgttttaacacggagtaagt gatgtaccgttatttgtctattgctgcggtggtactgagcgcagcattttccggcccggcgttggccgaaggtatcaatagtttt tctcaggcgaaagccgcggcggtaaaagtccacgctgacgcgcccggtacgttttattgcggatgtaaaattaactggcag ggcaaaaaaggcgttgttgatctgcaatcgtgcggctatcaggtgcgcaaaaatgaaaaccgcgccagccgcgtagagtg ggaacatgtcgttcccgcctggcagttcggtcaccagcgccagtgctggcaggacggtggacgtaaaaactgcgctaaa gatccggtctatcgcaagatggaaagcgatatgcataacctgcagccgtcagtcggtgaggtgaatggcgatcgcggcaa ctttatgtacagccagtggaatggcggtgaaggccagtacggtcaatgcgccatgaaggtcgatttcaaagaaaaagctgc cgaaccaccagcgcgtgcacgcggtgccattgcgcgcacctacttctatatgcgcgaccaatacaacc
  4.2 After editing ccggagccaaaactctcttacacccagcgcggaacctccgccggaacggcctggctggaaagctatgaaattcgcctcaa tcccgttttgctgttggaaaacagtgaagcttttattgaagaagtggtaccgcacgaactggcacatttgctggtatggaaaca tttcggccgcgtagcgccacatggcaaagagtggaagtggatgatggaaaacgtgctgggtgttcccgcccgtcgtacgc atcagttcgaactgcaatccgtgcgtcgcaacaccttcccctaccgctgcaagtgccaggagcatcagcttaccgtacgcc gccataatcgcgtagttcgtggcgaggccgtctatcgctgtgttcactgcggtgaacagctggttgcgaaataaccatctga actatcaggaactttcctgatctggctgattgcataccaaaacagctttcgctacgttgctggctcgttttaacacggagtaagt gatgtaccgttatttgtctattgctgATGTACCCCTTCTGTGTTGCcggtggtactgagcgcagcattttccg gcccggcgttggccgaaggtatcaatagtttttctcaggcgaaagccgcggcggtaaaagtccacgctgacgcgcccggt acgttttattgcggatgtaaaattaactggcagggcaaaaaaggcgttgttgatctgcaatcgtgcggctatcaggtgcgcaa aaatgaaaaccgcgccagccgcgtagagtgggaacatgtcgttcccgcctggcagttcggtcaccagcgccagtgctgg caggacggtggacgtaaaaactgcgctaaagatccggtctatcgcaagatggaaagcgatatgcataacctgcagccgtc agtcggtgaggtgaatggcgatcgcggcaactttatgtacagccagtggaatggcggtgaaggccagtacggtcaatgcg ccatgaaggtcgatttcaaagaaaaagctgccgaaccaccagcgcgtgcacgcggtgccattgcgcgcacctacttctata tgcgcgaccaatacaacc
5 *recA-*
  5.1 Before editing catcgcctggctcatcatacgtgccgcaaggcccatgtgagagtcgccgatttcgccttcgatttccgctttcggcgtcagtg ccgccacggagtcaacgacgataacgtctactgcgccagaacgcgccagggcgtcacagatttccagtgcctgctcgcc ggtgtccggctgggagcacagcaggttgtcgatatcgacgcccagtttacgtgcgtagattgggtccagcgcgtgttcagc atcgataaacgcacaggttttaccttcacgctgcgctgcggcgatcacctgcagcgtcagcgtggttttaccggaagattcc ggtccgtagatttcgacgatacggcccatcggcagaccacctgccccaagcgcgatatccagtgaaagcgaaccggtag agatggtttccacatccatggaacggtcttcacccaggcgcatgatggagcctttaccaaattgtttctcaatctggcccagtg ctgccgccaacgctttctgtttgttttcgtcgatagccatttttactcctgtcatgccgggtaataccggatagtcaatatgttctgt tgaagcaattatactgtatgctcatacagtatcaagtgttttgtagaaattgttgccacaaggtctgcaatgcatacgcagtagc ctgacgacgcaccgcatcacggtcgccgctgaagcattcccgccgggtaatgccttcaccgcgggcagtggcaaaagca aaccagacggtgccgacaggcttctcttcactgccgccatccggcccggcgataccactaatagacacggcataatcagc acgagccgctttcagtgcgcctatcgccatttccaccacgacgggttcactcaccgcgccatgctgcgccagcgtctcttcg cgtacgccgatcatctgcgctttggcttcgttactgtaggtgacaaatccgcgttcaaaccaggcggagctac
  5.2 After editing catcgcctggctcatcatacgtgccgcaaggcccatgtgagagtcgccgatttcgccttcgatttccgctttcggcgtcagtg ccgccacggagtcaacgacgataacgtctactgcgccagaacgcgccagggcgtcacagatttccagtgcctgctcgcc ggtgtccggctgggagcacagcaggttgtcgatatcgacgcccagtttacgtgcgtagattgggtccagcgcgtgttcagc atcgataaacgcacaggttttaccttcacgctgcgctgcggcgatcacctgcagcgtcagcgtggttttaccggaagattcc ggtccgtagatttcgacgatacggcccatcggcagaccacctgccccaagcgcgatatccagtgaaagcgaaccggtag agatggtttccacatccatggaacggtcttcacccaggcgcatgatggagcctttaccaaattgtttctcaatctggcccagtg ctgccgccaacgGGTCTTCTCATAGTATCGCCctttctgtttgttttcgtcgatagccatttttactcctgtcatg ccgggtaataccggatagtcaatatgttctgttgaagcaattatactgtatgctcatacagtatcaagtgttttgtagaaattgttg ccacaaggtctgcaatgcatacgcagtagcctgacgacgcaccgcatcacggtcgccgctgaagcattcccgccgggta atgccttcaccgcgggcagtggcaaaagcaaaccagacggtgccgacaggcttctcttcactgccgccatccggcccgg cgataccactaatagacacggcataatcagcacgagccgctttcagtgcgcctatcgccatttccaccacgacgggttcact caccgcgccatgctgcgccagcgtctcttcgcgtacgccgatcatctgcgctttggcttcgttactgtaggtgacaaatccgc gttcaaaccaggcggagctac

### Reference

1. Huo Y, Cho KM, Rivera JGL, Monte E, Shen CR, Yan Y, Liao JC: **Conversion of proteins into biofuels by engineering nitrogen flux.** *Nat Biotechnol* 2011, **29:**346-351.
2. Atsumi S, Hanai T, Liao JC: **Non-fermentative pathways for synthesis of branched-chain higher alcohols as biofuels.** *Nature* 2008, **451:**86-89.
